# Dopamine neurons drive basal forebrain reward prediction in milliseconds

**DOI:** 10.64898/2026.10.01.755874

**Authors:** You-Jhe Jhong, Ming-Ching Chiang, Adriane Guillaumin, Cheng-Yen Cheng, Mu-En Kuo, Cheng-Ze Lee, Alessandro Scaglione, Francois Georges, Shih-Chieh Lin

## Abstract

Phasic reward prediction signals in ventral tegmental area (VTA) dopamine (DA) neurons and basal forebrain (BF) noncholinergic bursting neurons are remarkably similar despite the distinct circuit organizations and functions of the two systems. Whether this similarity reflects independent computation within each system, inheritance from common upstream circuitry, or causal propagation from one system to the other remains unknown. Here we show that reward prediction signals are tightly coordinated across these two systems, with VTA DA activity causally driving BF bursting neurons on a millisecond timescale. Simultaneous recordings from freely moving rats revealed strong coupling between putative DA neurons and BF bursting neurons on individual trials, both in response amplitude and spike timing. DA activity consistently preceded BF bursting neuron activity by approximately 10 ms across well-trained behavior, learning, and even outside overt task events. In DAT-Cre mice, optogenetic activation of VTA DA neurons rapidly recruited BF bursting neurons at latencies matching those observed during natural DA-BF coupling, while activation of their BF terminals produced responses at still shorter latencies. Conversely, inhibition of DA neurons or their BF terminals suppressed both cue-evoked and, notably, ongoing activity in a substantial subset of BF bursting neurons. Fast DA-evoked BF recruitment persisted after D1/D2 receptor blockade, and anatomical tracing identified glutamatergic DA-to-BF synapses, supporting glutamate co-release as a candidate substrate. Together, these findings reveal a previously unrecognized causal link that enables rapid propagation of reward prediction signals between functionally distinct subcortical systems.

## Introduction

Phasic reward-prediction signals are expressed across multiple subcortical systems, notably in dopamine (DA) neurons in the ventral tegmental area (VTA)^1–6^ and a population of noncholinergic basal forebrain (BF) neurons termed BF bursting neurons^7–10^. These two populations exhibit remarkably similar reward-related activity, including rapid responses to reward-predicting cues whose amplitudes scale with expected value and change as cue–outcome associations are learned^1–10^. Yet they have been studied in largely separate functional and anatomical frameworks. VTA DA neurons are classically associated with reinforcement learning, value updating, and plasticity through their influence on striatal and limbic circuits^11–15^. In contrast, BF bursting neurons belong to a major corticopetal system that carries a gain-modulation signal linked to cortical activation, behavioral performance, and selective attention^7–9,16–18^. Thus, two neuronal populations studied in different behavioral contexts, assigned different functional roles, and projecting to largely distinct downstream circuits nevertheless carry strikingly similar reward-prediction signals. Why these signals should be so closely matched, and whether this similarity reflects any functional relationship between the two systems, remains unclear.

Several fundamentally different circuit organizations could account for this similarity. On the one hand, reward-prediction signals could be computed independently in the two systems to serve distinct functions. In this scenario, the DA and BF systems may have evolved to construct related quantities separately, allowing reward information to be used for learning and plasticity in one circuit and for behavioral performance or gain modulation in another. Their average response patterns could therefore appear similar even if their moment-to-moment fluctuations were largely independent. On the other hand, the two reward-prediction signals may be dynamically coordinated, such that their activity fluctuates together from moment to moment. Such coordination could reflect a shared upstream computation, in which a reward-prediction signal generated elsewhere is distributed in parallel to both systems^19^. Alternatively, it could reflect direct causal coupling, in which the signal is computed primarily in one system and propagated to the other through known anatomical projections^20–22^, linking circuits with different functional roles into a serial pathway. Thus, the similarity between DA and BF activity could arise from three fundamentally different organizations: independent computation within each system, parallel distribution of a common upstream computation, or causal propagation from one system to the other. Distinguishing among these architectures is therefore essential for understanding how reward-prediction signals are computed and distributed across the brain.

A key first step toward distinguishing among these architectures is to determine whether DA and BF reward-prediction signals are dynamically coupled from moment to moment, rather than merely resembling each other in their average response profiles. This requires recording the two populations simultaneously in the same animal under matched behavioral conditions, which we achieved in freely moving rats across reward-based behavioral contexts. Having established a strong coupling, the next question is whether the coordinated signals simply reflect common upstream drive or whether activity in one system causally influences the other. Because the many possible common upstream inputs to DA and BF cannot be exhaustively identified or excluded, we instead tested whether the temporally leading system exerts causal control over the lagging one using cell-type-specific manipulations of VTA DA neurons and their BF projections. Finally, given the unexpectedly rapid timescale of this coupling, we tested whether it depended on canonical dopamine receptor signaling^23–25^. These experiments establish a fast directional coupling from VTA DA neurons to BF bursting neurons that is coordinated on a trial-by-trial basis, robust across behavioral states, sufficient to drive BF bursting responses, necessary for a substantial population of these neurons, and independent of canonical DA-receptor signaling. Together, these findings show that a substantial component of the BF bursting reward-prediction signal is transmitted from the VTA DA system on a millisecond timescale, revealing direct causal coupling between two subcortical systems traditionally associated with distinct functions.

## Results

### Simultaneous recordings reveal that DA and BF response amplitudes are coupled on single trials

To determine whether VTA DA neurons and BF bursting neurons carry similar reward-prediction signals under identical behavioral conditions, we performed simultaneous dual-site electrophysiological recordings from the VTA and BF in freely moving rats (Figure 1A, Figure S1). Rats performed a lick-schedule reward task in which a trial-start signal indicated the availability of a new trial. Rats then initiated the trial by entering a fixation port, where one of three trial types occurred with equal probability (Figure 1B). In short-delay (SD) trials, the SD sound (S_SD_) signaled delivery of a water reward upon the third lick. In long-delay (LD) trials, a different sound (S_LD_) signaled delivery of the same reward between the 15th and 36th licks, depending on the session. In catch trials, both the sound and reward were omitted. Thus, SD and LD trials varied the number of required licks and the resulting delay to reward while holding reward magnitude constant. Rats behaviorally distinguished the two reward-predicting cues, producing shorter reaction times on SD than LD trials (0.296 ± 0.050s in SD trials vs 0.333 ± 0.048s in LD trials, mean ± s.e.m.; two-sided Wilcoxon signed-rank test, p<0.05) while maintaining comparable motivation to the two cues (SD: 99.63 ± 0.25%, LD: 98.94 ± 0.36% in hit rates; SD vs LD, mean ± s.e.m.; two-sided Wilcoxon signed-rank test with Bonferroni correction, p=0.9375) (Figure 1C). This behavioral difference is consistent with a higher expected value for the SD cue than for the LD cue, despite both predicting the same reward magnitude.

**Figure 1.**
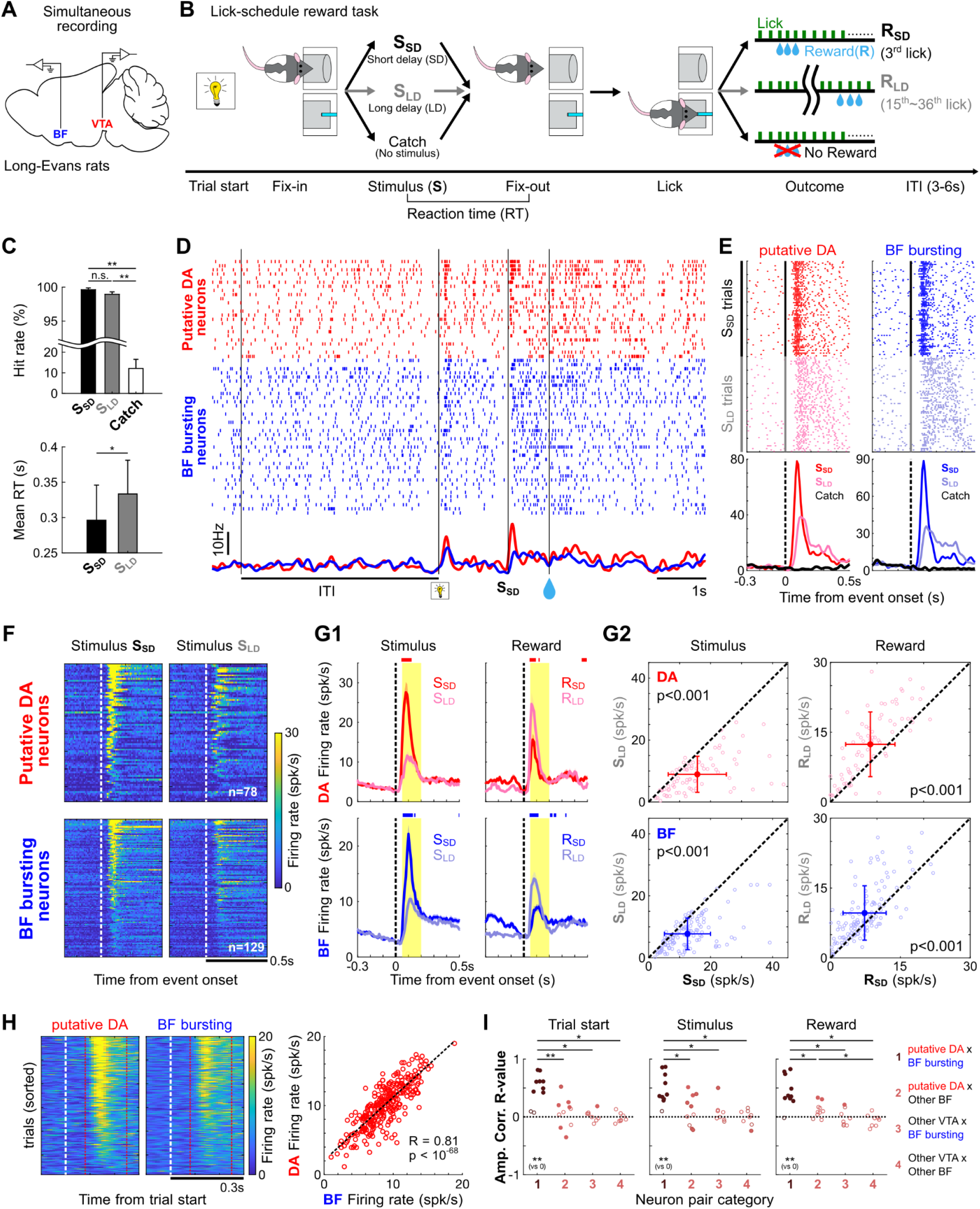
Putative DA neurons and BF bursting neurons show coupled response amplitudes on single trials. **A**, Schematic of simultaneous dual-site electrophysiological recordings from VTA and BF in freely moving Long-Evans rats. **B**, Lick-schedule reward task. Rats initiated each trial by entering the fixation port, after which one of three trial types occurred with equal probability. In short-delay (SD) and long-delay (LD) trials, distinct sounds (S_SD_ and S_LD_) predicted delivery of the same water reward after the third lick (R_SD_) or after 15-36 licks (R_LD_), respectively. Catch trials omitted both the sound and reward. Reaction time (RT) was measured from sound onset to fixation-port exit. **C**, Behavioral performance showing hit rate for each trial type (top) and mean RT on SD and LD trials (bottom). Data are mean ± s.e.m. across 10 recording sessions (from 7 rats, two-sided Wilcoxon signed-rank tests with Bonferroni correction. n.s., not significant; *p < 0.05; **p < 0.01.). **D**, Representative continuous recording segment showing spike rasters for all simultaneously recorded putative DA neurons (red) and BF bursting neurons (blue) within one session. Bottom, corresponding population firing rates averaged across neurons within each population. Task events are indicated below. **E**, Example putative DA neuron and BF bursting neuron from the same session showing similar modulation by S_SD_ and S_LD_ cues. **F**, Population heatmaps of responses to S_SD_ and S_LD_ for all putative DA neurons (n = 78) and BF bursting neurons (n = 129). Each row represents one neuron. Neurons were sorted by their average response amplitude 0.05-0.20 s after S_SD_ onset and are shown in the same order for S_LD_ responses. **G**, Similar coding of reward prediction values in putative DA and BF bursting neuron responses. **G1**, Population PSTHs aligned to sound onset and reward delivery on SD and LD trials. Traces show mean ± s.e.m. across neurons. Yellow shading indicates the response windows used for burst amplitude quantification. Colored horizontal bars indicate time bins with significant differences between SD and LD trials (two-sided Wilcoxon signed-rank tests with Bonferroni correction, p < 0.05). **G2**, Single-neuron responses to S_LD_ versus S_SD_ (left) and R_LD_ versus R_SD_ (right), quantified within the windows shown in G1. Open circles represent individual neurons, filled symbols and error bars indicate mean ± s.d., and dashed lines indicate equality. Response amplitudes differed significantly between the two trial types in all comparisons (two-sided Wilcoxon signed-rank tests, p < 0.001). **H**, Single-trial amplitude coupling in a representative session at the trial start event. Left, population firing-rate heatmaps for putative DA neurons and BF bursting neurons. Trials were sorted by putative DA response amplitude and are shown in the same order for BF bursting activity. Red lines indicate the response window used for quantification. Right, relationship between putative DA and BF bursting response amplitudes across individual trials. The dashed line indicates the linear fit; Pearson’s r and p value are shown. **I**, Trial-by-trial amplitude correlations across sessions at trial start, sound onset, and reward delivery. Pearson’s r values are shown for four population pairings: putative DA neurons with BF bursting neurons, putative DA neurons with other BF neurons, other VTA neurons with BF bursting neurons, and other VTA neurons with other BF neurons. Each circle represents one session (n=8-10 sessions); filled circles indicate significant within-session correlations (p < 0.01), and open circles indicate nonsignificant correlations. Each population pairing was compared with zero using a two-sided Wilcoxon signed-rank test with Bonferroni correction; only significant comparisons are indicated below each plot. Horizontal bars indicate significant differences between population pairings (one-sided Wilcoxon signed-rank tests with Bonferroni correction). n.s., not significant; *p < 0.05; **p < 0.01.

Simultaneous VTA–BF recordings captured activity from multiple putative DA neurons and BF bursting neurons within the same sessions (Figure 1D). These populations were operationally identified based on their phasic responses to reward-predicting cues (Figure S2), following established criteria^7–9,18,26^. Both populations showed larger responses to the higher-value SD cue than to the LD cue (two-sided Wilcoxon signed-rank tests, p<0.001) (Figure 1E-G). At reward delivery, this trial-type ordering was reversed: the same reward evoked smaller responses on SD than LD trials (two-sided Wilcoxon signed-rank tests, p<0.001) (Figure 1G). Thus, stronger responses to the higher-value cue were followed by weaker responses to the reward, consistent with the opposing modulation of cue- and reward-evoked activity by reward expectation in both dopamine neurons^27–30^ and BF bursting neurons^7,9,10^.

To determine whether these similar response patterns extended beyond trial averages, we used the simultaneous recordings to compare the response amplitudes of putative DA neurons and BF bursting neurons on individual trials within matched sessions and event types. In a representative session, trials with larger putative DA responses also showed larger BF bursting responses at trial start, producing a strong positive correlation (Figure 1H). Across animals and sessions, positive trial-by-trial correlations between the putative DA and BF bursting populations were consistently observed at trial start, stimulus, and reward (Pearson correlation of responses, p<0.01) (Figure 1I). Moreover, these correlations were significantly stronger than those observed in pairings involving other VTA or BF neurons (one-sided Wilcoxon signed-rank tests with Bonferroni correction, p<0.05) (Figure 1I; Figure S3), indicating selective coordination of response amplitudes between these populations. Together, these results argue against completely independent moment-to-moment activity and demonstrate selective coordination between putative DA neurons and BF bursting neurons.

### Millisecond spike-timing analyses reveal DA-leading BF coupling

Having shown that phasic response amplitudes were strongly coupled between putative DA neurons and BF bursting neurons (Figure 1I,J; Figure S3), we next asked whether their activity was also coordinated in time. Raw spike rasters from all identified neurons in a representative session (Figure 1D) suggested that this coordination extended beyond discrete phasic events. Ongoing fluctuations in the two populations appeared closely matched across the session, including periods outside task-evoked bursts. This observation motivated direct analysis of their spike-timing relationship at millisecond resolution.

We computed normalized spike-time cross-correlation functions at 1-ms temporal resolution between simultaneously recorded VTA–BF neuron pairs, using all spikes recorded throughout each session (Figure 2A, Figure S4). Putative DA neuron-BF bursting neuron pairs showed strong correlations near zero lag (Figure 2A), which were not observed for other VTA–BF pairings (Figure S4). With putative DA spikes as the reference, the overall mean cross-correlation function peaked at a 10-ms DA-leading lag (Figure 2A). This temporal asymmetry was consistent across all five animals: among significantly correlated pairs, the interquartile range of peak lags lay entirely in the DA-leading direction in each animal. Restricting the analysis to spikes recorded during intertrial intervals preserved both the correlation and the approximately 10-ms DA-leading offset (Figure 2B), indicating that the temporal asymmetry was not generated solely by shared locking to overt task events.

**Figure 2.**
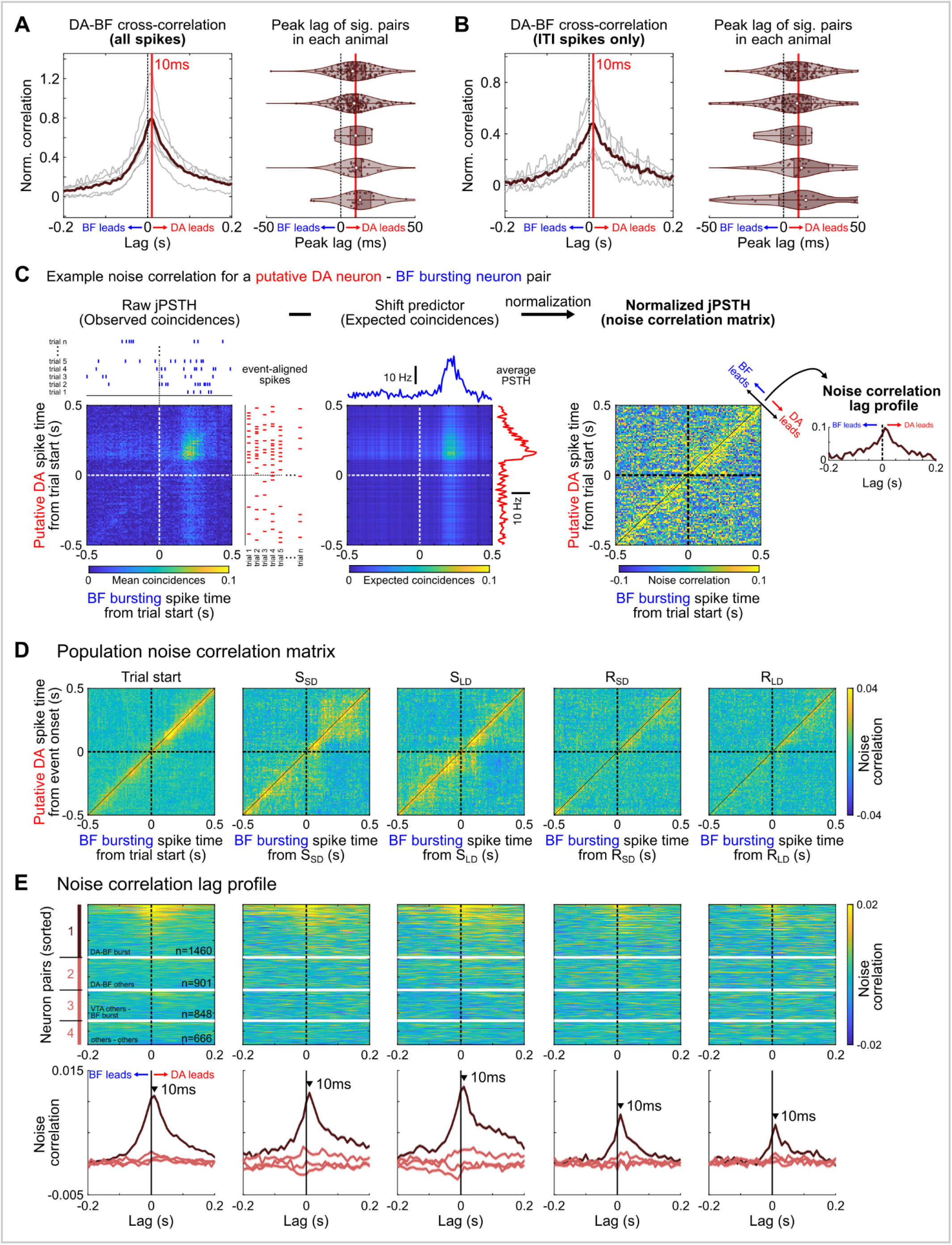
Putative DA activity precedes BF bursting activity by approximately 10 ms. **A,** Normalized spike-time cross-correlation analysis using all spikes recorded throughout each behavioral session. Cross-correlograms were normalized as (observed − expected)/expected, expressing the relative excess in coincidence counts rather than a Pearson correlation coefficient. Zero indicates the expected count, and a value of 1 indicates twice that count. Left, average cross-correlation functions for significantly correlated putative DA neuron-BF bursting neuron pairs in each animal (gray; n = 5 rats) and the overall mean across animals (brown). The red line marks the peak of the overall mean function, with putative DA activity temporally leading BF bursting activity by 10 ms. Right, peak-lag distributions for all significantly correlated neuron pairs, grouped by animal. Open white circles indicate medians, and darker shading indicates the interquartile range (25th-75th percentiles). **B**, The same spike-time cross-correlation analysis restricted to spikes recorded during intertrial intervals (ITIs). Conventions are as in **A**. The persistence of the DA-leading peak during ITIs suggests that the temporal relationship was not generated solely by shared responses to task events. **C**, Schematic of the noise correlation analysis for an example putative DA neuron-BF bursting neuron pair, shown for activity aligned to the trial start event. The normalized jPSTH, termed the noise correlation matrix, was calculated in 10-ms bins by correcting the raw jPSTH for expected event-locked covariation and normalizing by firing-rate variability. Horizontal and vertical dashed lines indicate event onset, and the solid diagonal indicates zero lag. Averaging along matrix diagonals representing constant time lags produced a one-dimensional noise correlation lag profile. **D**, Population noise correlation matrices for the same significantly correlated putative DA neuron-BF bursting neuron pairs included in **A**, at trial start, SD and LD stimulus onset, and SD and LD reward delivery. **E**, Noise correlation lag profiles for all simultaneously recorded neuron pairs. Top, individual lag profiles grouped into the four population-pairing categories defined in Figure 1I. Each row represents one neuron pair. Bottom, mean lag profile (± s.e.m.) for each pairing category. Vertical black lines indicate zero lag, and arrowheads mark the peak for each event condition. Only putative DA neuron-BF bursting neuron pairs exhibited a prominent DA-leading peak across all event conditions, demonstrating the selectivity of this lagged temporal relationship.

We next asked whether this temporal relationship represented a persistent feature of their activity or arose primarily from phasic responses to shared task events. To isolate coupling beyond shared event-locked activity, we performed noise-correlation analyses using 10-ms bins, removing the mean event-locked response for each condition (Figure 2C). Across key task epochs, including trial start, cue, and reward, the residual correlations consistently exhibited a DA-leading displacement (Figure 2D), with lag profiles peaking at 10 ms, corresponding to a one-bin offset at this temporal resolution (Figure 2E). Moreover, this displacement persisted throughout each event-aligned analysis window, extending beyond the phasic response itself (Figure 2D). This residual temporal coupling was selective for putative DA and BF bursting neurons and was not observed for other VTA–BF neuronal pairings (Figure 2E). Together, these analyses demonstrate a consistent fine-timescale relationship across ongoing and task-related activity, in which putative DA activity precedes BF bursting activity by approximately 10 ms.

### DA-leading coupling persists throughout reward learning

One potential concern was that the amplitude and timing coupling described above had thus far been established only in animals performing a well-learned, overtrained task. Under these conditions, stable reward-prediction information could be redundantly represented across multiple brain regions, leaving open the possibility that matched DA and BF activity emerged only after extensive training or that their relationship might break down while reward predictions were being updated. To test these possibilities, we recorded from a separate cohort of rats across three consecutive learning phases in which cue-reward relationships were acquired, extinguished, or degraded (Figure 3A). Throughout all phases, a familiar, well-learned auditory cue continued to predict reward, providing an internal control and allowing putative DA neurons and BF bursting neurons to be identified using the same functional criteria across sessions. Rats first learned that a novel visual cue also predicted reward, then underwent extinction of this newly acquired association, and finally experienced contingency degradation in which additional rewards were delivered following spontaneous licking, independently of cue presentation. For each learning phase, we analyzed the session showing the largest within-session behavioral change.

**Figure 3.**
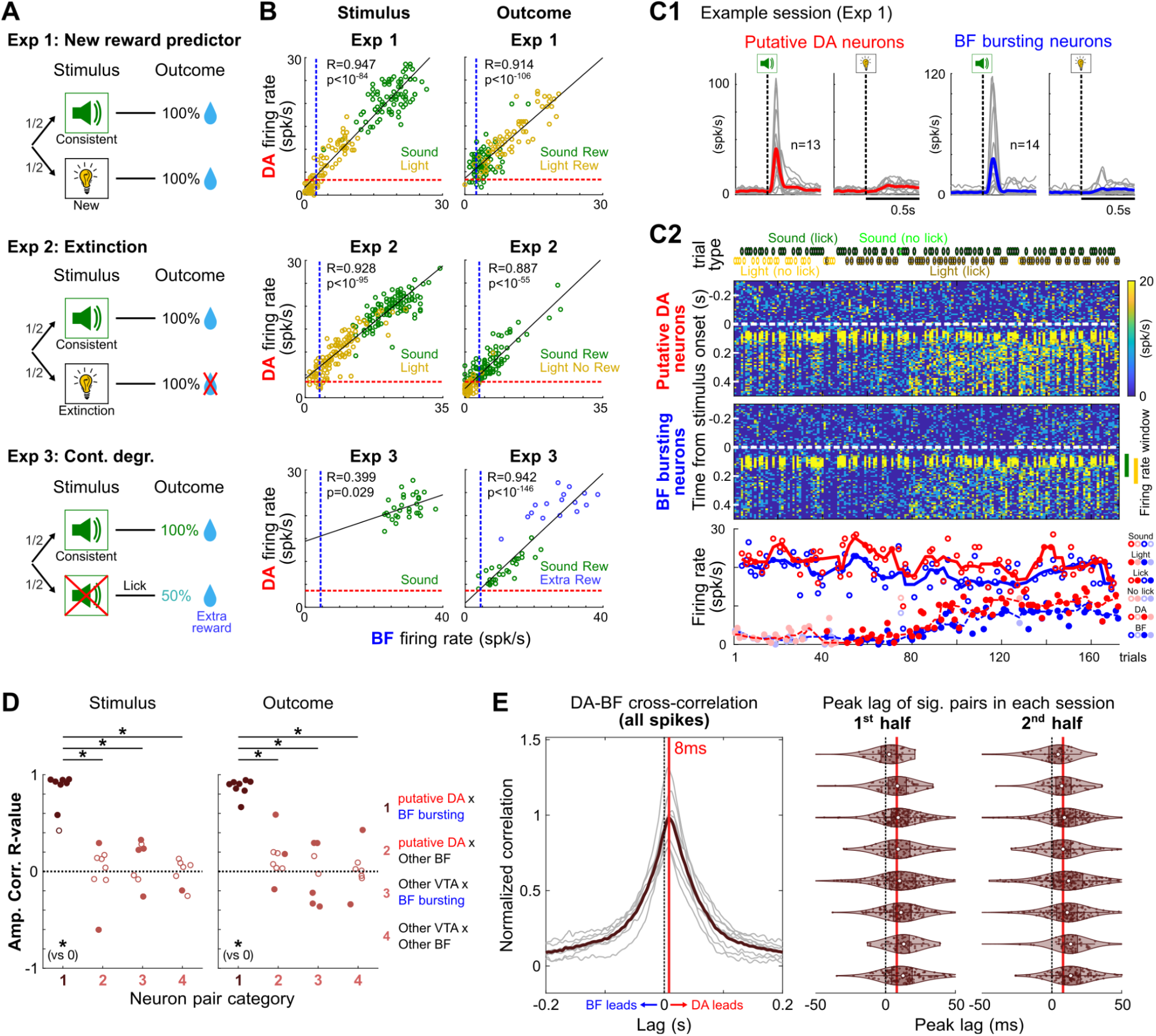
DA-leading coupling persists throughout reward learning. **A**, Sequence of three learning paradigms in freely moving rats: acquisition of a new light-reward association, extinction, and contingency degradation. A familiar sound remained consistently paired with reward throughout all phases and served as an internal control. **B**, Trial-by-trial amplitude coupling in example sessions from Experiments 1–3, shown from top to bottom. Putative DA and BF bursting population firing rates are shown during the stimulus (left) and outcome (right) epochs. Open circles represent individual trials, colors indicate trial conditions, and black lines indicate linear fits. Blue and red dashed lines indicate the baseline firing rates of the BF bursting and putative DA populations, respectively. **C**, Evolution of putative DA and BF bursting activity during an example learning session from Experiment 1. **C1**, Putative DA neurons and BF bursting neurons were identified by their phasic responses to the consistent sound. Sound- and light-aligned PSTHs are shown for simultaneously recorded putative DA neurons (red; n = 13) and BF bursting neurons (blue; n = 14). Gray traces represent individual neurons, and colored traces represent population averages. **C2**, Trial-by-trial development of stimulus responses in the same session. Trial types and licking outcomes are indicated above the heatmaps. Heatmaps show stimulus-aligned population firing rates across successive trials for putative DA neurons (top) and BF bursting neurons (middle), with each column representing one trial. White dashed lines indicate stimulus onset. The bars to the right of the lower heatmap mark the analysis window used to calculate the response amplitudes shown below. Bottom, corresponding response amplitudes across trials, with symbols and colors defined by the key at right. Lines show moving medians across learning. Despite substantial trial-to-trial variation in activity, DA and BF population dynamics remained closely matched throughout learning. **D**, Session-wise Pearson correlations of response amplitudes between simultaneously recorded neuronal populations during the stimulus (left) and outcome (right) epochs. Conventions are as in Figure 1I. Horizontal bars indicate significant differences between population pairings (one-sided Wilcoxon signed-rank tests with Bonferroni correction, *p < 0.05). Consistent with Figure 1I, strong positive response-amplitude correlations were selective for putative DA neuron-BF bursting neuron pairs. **E**, Fine temporal coupling across learning contexts. Left, cross-correlation functions for significantly correlated putative DA neuron-BF bursting neuron pairs. Gray lines represent individual session averages, and the overall mean peaked at an 8-ms DA-leading lag. Right, peak-lag distributions for each session, calculated separately using spikes from the first and second halves of the session. The DA-leading relationship persisted in both halves. Other conventions are as in Figure 2A.

Putative DA and BF bursting responses continued to change together as reward predictions were updated. Example sessions from each learning phase showed coordinated trial-by-trial variation in stimulus- and outcome-related response amplitudes (Pearson correlation) (Figure 3B). In the acquisition example, both populations retained strong responses to the familiar sound used for functional identification (Figure 3C1). As responses to the novel light cue changed during learning, response amplitudes in the two populations remained closely coupled on individual trials throughout the session (Figure 3C2). DA and BF activity was similarly matched during extinction and contingency degradation (Figure S5). Across sessions, trial-by-trial stimulus- and outcome-related response amplitudes were strongly correlated between the putative DA and BF bursting populations, with correlations significantly exceeding those observed for other VTA–BF population pairings (one-sided Wilcoxon signed-rank tests with Bonferroni correction, p<0.05) (Figure 3D). Thus, selective amplitude coupling persisted as reward predictions were updated.

Spike-time cross-correlation analyses next tested whether the precise temporal relationship between the two populations was maintained during learning. Across all three learning phases, the mean cross-correlation function retained an 8-ms DA-leading peak (Figure 3E). To determine whether this timing relationship also remained stable within individual sessions, we computed cross-correlations separately using spikes from the first and second halves of each session. Peak-lag distributions for significantly correlated pairs remained consistently DA-leading in both halves, with the interquartile range lying almost entirely in the DA-leading direction in each session (Figure 3E). Thus, putative DA neurons and BF bursting neurons remained coupled in both response amplitude and spike timing as reward predictions were acquired, extinguished, or degraded, with putative DA activity preceding BF bursting activity by approximately 10 ms.

### Activation of DA neurons and their axon terminals rapidly evokes spiking in BF bursting neurons

Our results thus far have established that putative DA neurons and BF bursting neurons are closely coordinated in both response amplitude and spike timing, arguing against independent computation of reward prediction signals in the two populations. However, the consistent DA-leading coupling remains compatible with two circuit architectures. In a lagged common-input model, a reward prediction signal is generated by a common upstream source and reaches DA neurons before BF bursting neurons, allowing the two populations to operate in parallel without directly interacting. Alternatively, DA neurons may themselves drive BF bursting activity through direct causal coupling. We reasoned that exhaustively testing the lagged common-input scenario would be impractical given the many potential upstream sources that could influence the two populations through direct or indirect pathways^19,31–33^. In contrast, direct causal coupling makes experimentally testable predictions: selective activation of DA neurons should be sufficient to evoke BF bursting, whereas DA inhibition should suppress it.

We first tested the sufficiency prediction using dopamine transporter-Cre (DAT-Cre) mice to express channelrhodopsin-2 (ChR2) selectively in genetically defined VTA DA neurons (Figures 4A, S6). We recorded BF activity in head-fixed mice performing an auditory cue-reward association task, allowing BF bursting neurons to be identified by their phasic responses to the reward-predicting cue using the same operational criterion applied in rats. We subsequently activated VTA DA neurons outside the behavioral task, in the absence of sensory cues or reward, thereby isolating BF responses to DA activation from activity evoked by shared behavioral events.

**Figure 4.**
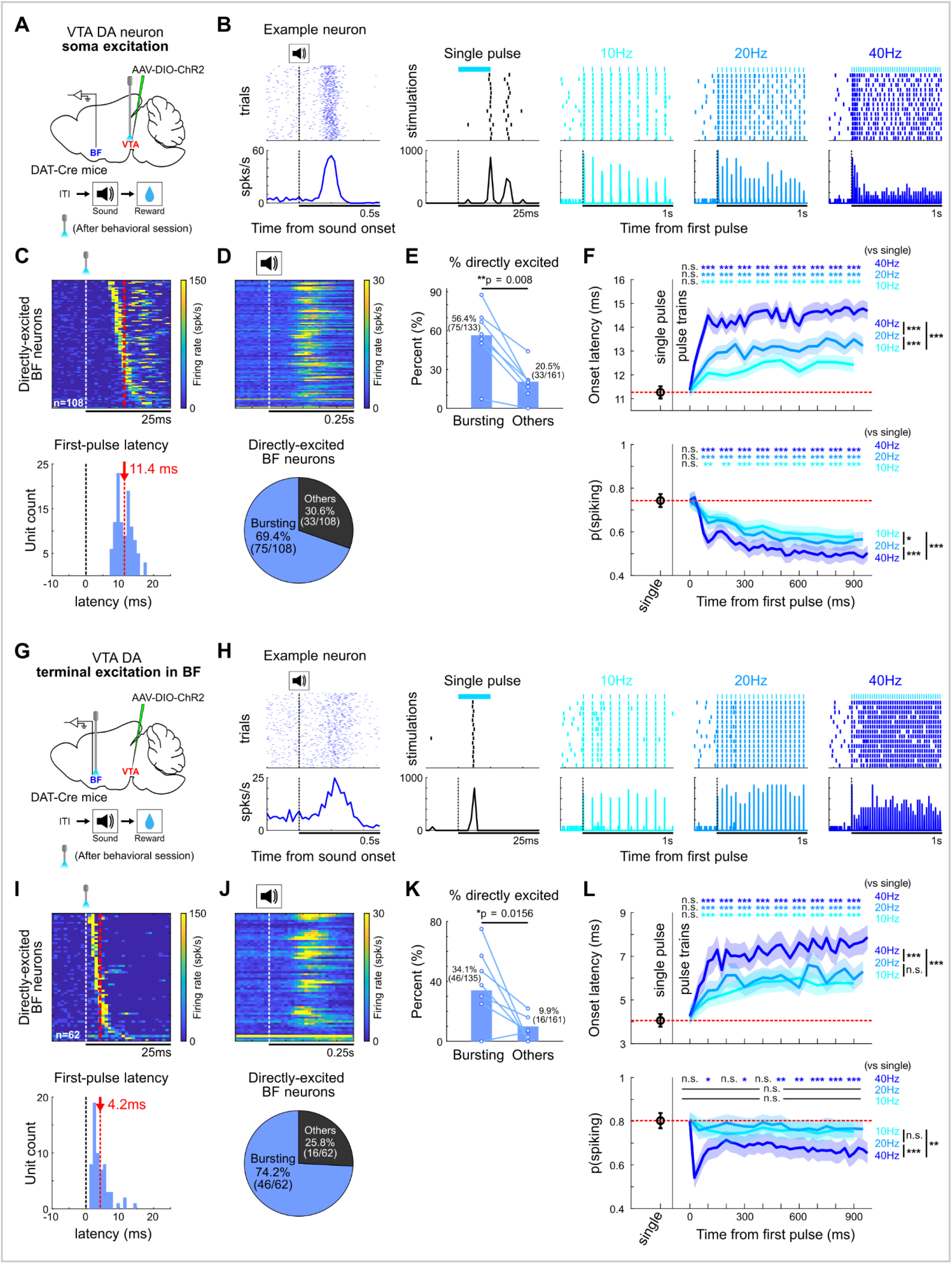
Activation of DA neurons and their axon terminals rapidly evokes spiking in BF bursting neurons. **A**, Somatic DA activation experiment. ChR2 was expressed in VTA DA neurons of DAT-Cre mice, and BF neurons were recorded during an auditory cue-reward task and subsequent optogenetic stimulation outside the task. **B**, Responses of an example BF bursting neuron to the reward-predicting sound and to a single 10-ms light pulse or 1-s stimulation trains at 10, 20, or 40 Hz. Light-pulse timing is shown above the spike rasters. **C**, Light-evoked activity of directly excited BF neurons, sorted by first-pulse latency (top), and distribution of first-pulse latencies (bottom; mean, 11.4 ms). Each row represents one neuron. **D**, Sound-evoked activity of the same neurons in the same order (top) and composition of the directly excited population (bottom). Most were BF bursting neurons (75/108; 69.4%). **E**, Proportions of BF bursting neurons and other BF neurons that were directly excited. Circles indicate individual mice (n=7), with paired measurements connected (one-sided Wilcoxon signed rank test). **F**, Onset latency (top) and spiking probability (bottom) following a single pulse and successive pulses within 10-, 20-, and 40-Hz trains. Red dashed lines indicate the single-pulse response levels for reference; colored traces show mean ± s.e.m. Asterisks indicate comparisons with single-pulse responses (two sided Wilcoxon signed rank test with Bonferroni correction). Comparison between train-pulse responses used a one-way repeated measures ANOVA with Bonferroni correction. n.s., not significant; *p < 0.05; **p < 0.01; ***p < 0.001. **G-L**, Corresponding experiment and analyses during activation of VTA DA terminals in the BF. Conventions are as in **A-F**. The mean first-pulse latency was 4.2 ms (**I**), and most directly excited neurons were BF bursting neurons (46/62; 74.2%; **J**).

Brief somatic activation of VTA DA neurons with 10-ms, 465-nm light pulses (4–6 mW) produced reliable, time-locked spiking responses in a subset of BF neurons (Figure 4B). Across all recorded BF neurons, 36.7% (108/294) exhibited short-latency excitation following somatic DA neuron activation (Figure S7), and the mean latency of BF responses to the first stimulation pulse was 11.4 ms (Figure 4C), closely matching the approximately 10-ms DA-leading offset observed during simultaneous recordings in freely moving rats. Among these directly excited neurons, the majority (69.4%) were BF bursting neurons (Figure 4D). The proportion of directly excited neurons was 56.4% among BF bursting neurons, significantly higher than 20.5% among other BF neurons (one-sided Wilcoxon signed rank test, p=0.016) (Figure 4E). Moreover, successive pulses within a stimulation train evoked BF responses with longer latencies and lower spiking probabilities compared with single-pulse responses (two sided Wilcoxon signed rank test with Bonferroni correction) (Figure 4F), consistent with a synaptically mediated process. Somatic DA activation therefore reproduced both the population selectivity and temporal delay of the naturally occurring DA-BF coupling, linking the optogenetically recruited neurons to the same operationally defined BF population studied in rats.

Somatic activation alone could not determine whether BF recruitment was mediated by DA axons within the BF or indirectly through other DA targets. We therefore stimulated DA axon terminals within the BF while recording nearby BF neurons (Figure 4G). Terminal stimulation reliably evoked time-locked spiking responses in BF neurons (Figure 4H). Across all recorded BF neurons, 20.9% (62/296) exhibited short-latency excitation (Figure S7). Critically, the mean response latency following terminal stimulation was 4.2 ms, substantially shorter than that following somatic stimulation (Figure 4I), consistent with bypassing propagation from DA somata to their BF axon terminals. Other response properties mirrored those observed with somatic activation, including preferential recruitment of BF bursting neurons and progressively longer latencies and lower spiking probabilities across successive pulses (Figure 4J-L). These findings identify DA axons within the BF as a local pathway sufficient for the fast recruitment of BF bursting neurons.

Together, these experiments establish that activation of VTA DA neurons or their axon terminals within the BF is sufficient to rapidly recruit BF bursting neurons. Somatic activation reproduced the approximately 10-ms delay observed during behavior, whereas terminal activation evoked responses at still shorter latencies.

These findings are notable because, although dopaminergic projections from the VTA to the VP have been anatomically documented^20^, DA innervation of the substantia innominata and adjacent posterior VP appears relatively sparse compared with that of the striatum^22^. Moreover, the latency of BF recruitment is substantially faster than expected from canonical G-protein-coupled dopamine receptor signaling^23–25^. These results therefore strongly constrain the potential mechanisms underlying the rapid DA-to-BF interaction.

### Inhibition of DA neurons and their axon terminals rapidly suppresses BF bursting-neuron activity

Having established that DA neuron activation is sufficient to evoke spiking in BF bursting neurons, we next tested whether DA activity is also required to sustain their activity. If DA-to-BF coupling conveys only phasic reward-prediction signals, inhibiting DA neurons should reduce cue-evoked bursting while sparing ongoing BF activity. Alternatively, if DA neurons provide continuous excitatory drive, inhibition should also suppress BF firing in the absence of a sensory event. We therefore inhibited DA neurons on a randomly selected half of sound and catch trials, allowing within-session comparisons with non-inhibited trials of each type (Figure 5A). Sound trials tested the contribution of DA activity to cue-evoked bursting, whereas catch trials tested its contribution to ongoing firing.

**Figure 5.**
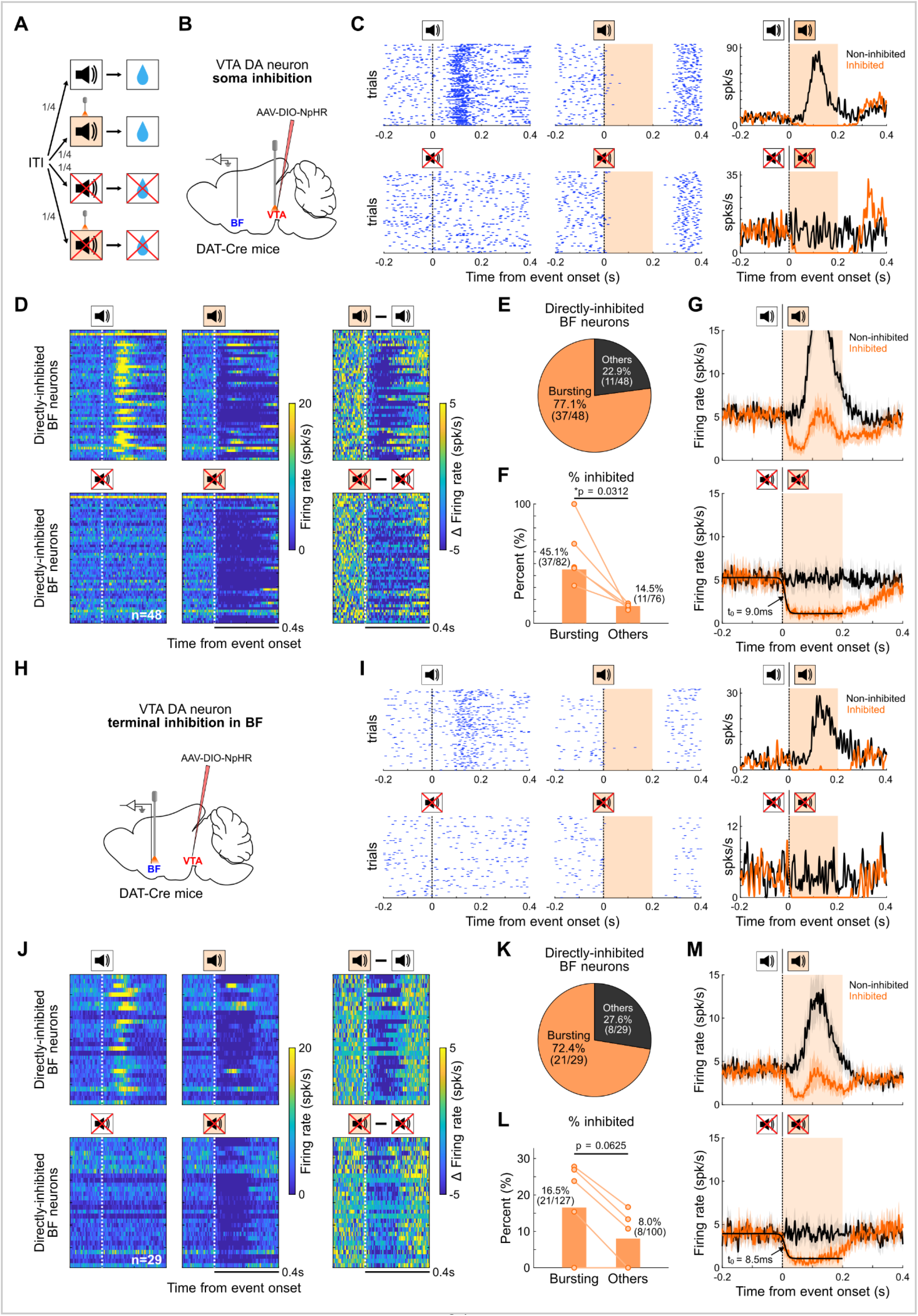
Inhibition of DA neurons and their axon terminals rapidly suppresses BF bursting-neuron activity. **A**, Trial structure. DA inhibition was delivered on a randomly selected half of sound and catch trials, allowing within-session comparisons with non-inhibited trials of each type. **B**, Experimental design for somatic DA inhibition. eNpHR3.0 was expressed in VTA DA neurons of DAT-Cre mice, and BF activity was recorded during optogenetic inhibition of DA somata. **C**, Example BF bursting neuron in which somatic DA inhibition nearly abolished both cue-evoked spiking on sound trials (top) and ongoing firing on catch trials (bottom). Rasters and PSTHs are shown for non-inhibited (black) and inhibited (orange) trials. Orange shading indicates the inhibition period. **D**, Population responses of directly inhibited BF neurons on sound (top) and catch (bottom) trials. Heatmaps show activity on non-inhibited and inhibited trials and the difference between conditions. Each row represents one neuron. Inhibition lasted 200 ms in most sessions and 500 ms in a subset of sessions on both sound and catch trials. **E**, Composition of the directly inhibited population. Most directly inhibited neurons were BF bursting neurons (37/48; 77.1%). **F**, Proportions of directly inhibited neurons among BF bursting and other BF neurons. Circles indicate individual animals (n=5), with measurements from the same animal connected. Direct inhibition was more frequent among BF bursting neurons (one-sided Wilcoxon signed rank test). **G**, Population responses of directly inhibited neurons on sound (top) and catch (bottom) trials, with and without somatic DA inhibition. A sigmoid fit to the inhibited catch-trial response yielded an estimated inhibition onset of 9.0 ms after light onset. **H–M**, Corresponding DA axon-terminal inhibition experiment. **H**, Experimental design; **I**, example BF bursting neuron; **J**, population responses; **K**, composition of the directly inhibited population; **L**, proportions of directly inhibited neurons among BF bursting and other BF neurons; **M**, population estimate of inhibition onset. Conventions are as in **B–G**. Most directly inhibited neurons were BF bursting neurons (21/29; 72.4%), and direct inhibition affected 16.5% of BF bursting neurons and 8.0% of other BF neurons. A sigmoid fit to the inhibited catch-trial response yielded an estimated inhibition onset of 8.5 ms.

To selectively inhibit VTA DA neurons, we expressed halorhodopsin (eNpHR3.0) in DAT-Cre mice and recorded BF activity during optogenetic inhibition of DA somata (Figures 5B, S6). We found that, in many cases, DA somatic inhibition not only nearly abolished the cue-evoked burst on sound trials but also rapidly and nearly completely suppressed ongoing activity on catch trials, as illustrated by the representative neuron (Figure 5C). Approximately 30% (48/158) of recorded BF neurons were classified as directly inhibited based on a significant reduction in firing within 25 ms after light onset (Figure S8). Population heatmaps revealed rapid suppression on both trial types and near-complete elimination of ongoing catch-trial activity in many neurons (Figure 5D). The vast majority (77.1%) of directly inhibited neurons were BF bursting neurons (Figure 5E). The proportion of directly inhibited neurons was 45.1% among BF bursting neurons, significantly higher than 14.5% among other BF neurons (one-sided Wilcoxon signed rank test, p=0.0312) (Figure 5F). To estimate the onset of inhibition, we pooled directly inhibited neurons on catch trials, where suppression was nearly complete, to improve the reliability of the estimate given sparse baseline spiking over short windows. Because small absolute changes in firing rate can delay significance-based detection of inhibition onset, we instead fitted a sigmoid to the population response to estimate the transition point. The fitted transition point occurred 9.0 ms after light onset (Figure 5G), placing somatic inhibition on the same approximately 10-ms timescale as the natural DA-leading relationship. The near-complete suppression of catch-trial activity demonstrates that DA neurons provide ongoing excitatory drive to the affected BF bursting neurons rather than contributing only to their phasic cue responses.

Having established through ChR2-mediated terminal activation that DA projections within the BF are sufficient to evoke BF spiking, we next inhibited the same projection to test whether its activity is required to sustain BF firing (Figure 5H). Terminal inhibition reproduced the key effects of somatic inhibition, rapidly suppressing both cue-evoked bursting and ongoing catch-trial activity in a representative neuron (Figure 5I) and across the directly inhibited population (Figure 5J). Overall, terminal inhibition directly affected 12.8% (29/227) of recorded BF neurons (Figure S8). As with somatic inhibition, most directly inhibited neurons (72.4%) were BF bursting neurons (Figure 5K). The proportion of directly inhibited neurons was 16.5% among BF bursting neurons, compared with 8.0% among other BF neurons (one-sided Wilcoxon signed rank test, p=0.0625) (Figure 5L). The relatively small number of directly inhibited neurons and small absolute changes in firing rate limited the precision of estimates of inhibition onset. Nevertheless, the same pooled catch-trial analysis yielded an estimated latency of approximately 8.5 ms (Figure 5M), placing terminal-mediated suppression on the same approximately 10-ms timescale as somatic inhibition. Together with the activation results, these findings show that local DA projections can rapidly evoke BF spiking and are required to sustain both cue-evoked and ongoing activity in a subset of BF bursting neurons.

### Fast DA-to-BF coupling persists during D1/D2 receptor blockade

The approximately 10-ms timing of DA-to-BF coupling is substantially faster than canonical D1 and D2 receptor signaling, which operates through G-protein-coupled second-messenger cascades^23–25^. This temporal mismatch raised the question of whether the fast recruitment of BF neurons requires canonical DA receptor signaling.

To test this, we measured BF responses to optogenetic activation of VTA DA somata before and every 10 min for 60 min after systemic administration of the D1 and D2 receptor antagonists SCH23390 and raclopride, or PBS as a control (Figures 6A, S6). The antagonists were administered at doses previously shown to alter striatal firing and local field potential oscillations^34^. In our cohort, antagonist administration also altered BF LFP activity and locomotion (Figure S9). Despite these effects, somatic DA activation continued to evoke short-latency BF responses for up to 60 min, as illustrated by a representative neuron (Figure 6B). Across the population, 69 of 77 neurons responsive before injection retained responses during both post-injection intervals (10–30 and 40–60 min; Figure 6C1,C2). A similar pattern was observed in PBS controls, with 65 of 70 initially responsive neurons retaining responses during both intervals (Figure 6D1,D2). In both the antagonist and PBS groups, persistently responsive neurons showed the most robust responses to DA stimulation, whereas neurons whose responses disappeared or emerged after injection showed much weaker responses (Figure 6C1,D1).

**Figure 6.**
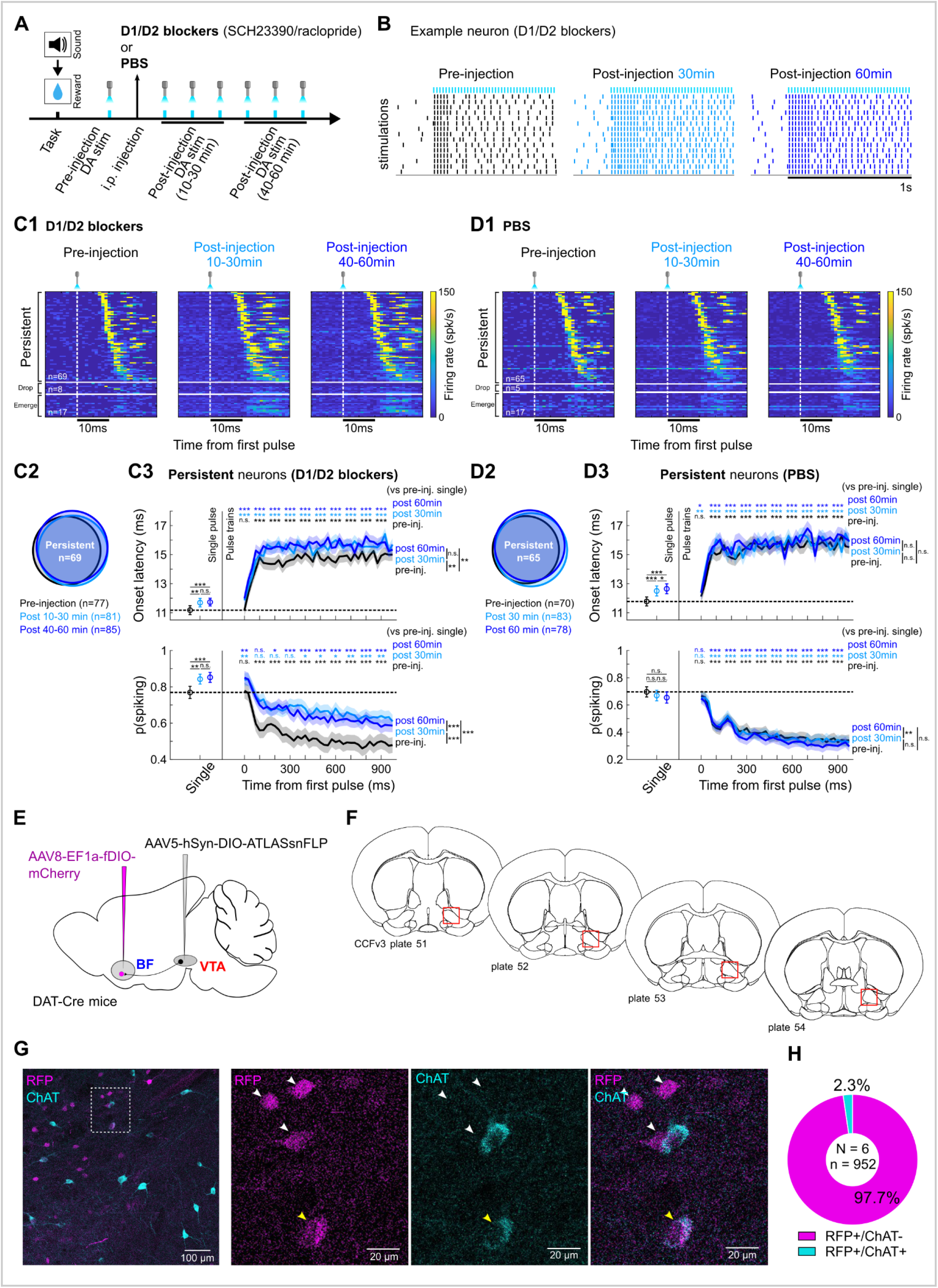
Fast DA-to-BF coupling does not depend on D1/D2 receptor signaling and is anatomically consistent with glutamate co-release. **A**, Experimental timeline. After behavioral characterization, BF responses to somatic DA stimulation were measured before and every 10 min for 60 min after systemic injection of D1/D2 antagonists (SCH23390/raclopride) or PBS. Analyses compared pre-injection (black), 10–30 min (light blue), and 40–60 min (dark blue) post-injection responses. **B**, Responses of an example persistently responsive BF neuron to DA stimulation before and after D1/D2 receptor blockade. Colored ticks indicate light pulses. **C**, BF responses before and after D1/D2 receptor blockade. **C1**, Heatmaps of responses to DA stimulation, grouped according to whether responses persisted, disappeared, or emerged after injection. Each row represents one neuron, with the same neuronal order across measurement periods. **C2**, Overlap among directly excited BF neurons across the three measurement periods. **C3**, Response onset latency (top) and spiking probability (bottom) in persistently responsive neurons following a single light pulse and successive pulses within stimulation trains. Traces show mean ± s.e.m. Asterisks above the traces indicate comparisons with the pre-injection single-pulse response (two-sided Wilcoxon signed-rank tests); brackets indicate pairwise comparisons between measurement periods following one-way repeated-measures ANOVA. All pairwise comparisons were Bonferroni-corrected. n.s., not significant; *p < 0.05; **p < 0.01; ***p < 0.001. Fast BF recruitment persisted after combined D1/D2 receptor blockade. **D**, Corresponding analyses following PBS administration. Conventions are as in **C**. **E**, Viral approach using the ATLAS strategy to label BF neurons postsynaptic to VTA DA axons at GluA1-containing glutamatergic synapses in DAT-Cre mice. **F**, Schematic representation of the coronal brain levels used to quantify RFP+ neurons in the BF. **G**, Representative confocal image of the RFP+ and ChAT+ neurons in the BF with close-up images. White arrowheads indicate RFP+/ChAT- neurons and the yellow arrowhead a RFP+/ChAT+ neuron. ChAT (choline acetyltransferase) labels BF cholinergic neurons. **H**, Quantification of RFP+/ChAT- and RFP+/ChAT+ neurons in the BF. 97.7% of ATLAS labeled neurons were ChAT-negative.

Among persistently responsive neurons, first-pulse response latencies increased slightly but significantly after antagonist administration, with a similar increase following PBS injection (Figure 6C3,D3). Successive stimulation pulses continued to evoke longer latencies and lower spiking probabilities in both groups. Spiking probability increased after D1/D2 receptor blockade but not after PBS injection, yet DA stimulation continued to evoke short-latency BF responses (Figure 6C3,D3). Thus, although response latency and spiking probability changed after injection, fast optogenetically evoked BF recruitment persisted despite D1/D2 receptor blockade.

Finally, we sought anatomical evidence for a possible fast synaptic mechanism. Glutamate co-release from VTA DA neurons has been demonstrated at other projection targets, particularly the ventral striatum^35–37^. We leveraged ATLAS (Anterograde Transsynaptic Label based on Antibody-like Sensors)^38^, a recently developed monosynaptic transsynaptic tracer that labels postsynaptic neurons through GluA1-containing glutamatergic synapses, to test whether VTA DA axons form glutamatergic connections in the BF and to identify the neurons they target. We expressed Cre-dependent ATLAS-FLP in VTA DA neurons and a FLP-dependent mCherry reporter locally within the BF of DAT-Cre mice (Figure 6E). ATLAS-labeled postsynaptic neurons were identified across the BF region examined (Figure 6F,G).

ChAT immunostaining revealed that 97.7% of labeled neurons were ChAT-negative, whereas only 2.3% were ChAT-positive (n = 952 neurons from N = 6 mice; Figure 6H). VTA DA neurons therefore establish glutamatergic synaptic contacts with a predominantly noncholinergic BF population. Together, the convergence of pharmacology that rules out canonical DA receptor signaling with anatomy demonstrating glutamatergic synapses from DA terminals onto noncholinergic BF neurons supports glutamate co-release as a leading candidate mechanism for fast DA-to-BF coupling.

## Discussion

Our findings identify fast causal coupling between VTA DA neurons and BF bursting neurons as a link between their reward-prediction signals. Simultaneous recordings revealed coupled response amplitudes and a consistent approximately 10-ms DA-leading relationship that persisted during learning and outside overt task events (Figures 1–3). Somatic and terminal manipulations established that DA neurons rapidly influence both cue-evoked and ongoing BF activity (Figures 4,5). The persistence of fast recruitment during D1/D2 receptor blockade, together with anatomical evidence for glutamatergic connections onto noncholinergic BF neurons, supports glutamate cotransmission as a candidate substrate (Figure 6).

### Fast causal coupling links DA and BF reward-prediction activity

The principal conceptual advance of the current study is to recast DA and BF reward-prediction signals from analogous but potentially separate representations into components of a directionally coupled circuit. Previous observations that both populations respond similarly to reward-predicting cues and outcomes could not distinguish independent but convergent computation from common upstream control or causal coupling^1–10^. The combination of amplitude coupling, directional timing, and causal manipulation now supports an organizational model in which DA activity contributes directly to the generation of BF reward-prediction activity.

The existence of DA-to-BF coupling means that a lagged common input is not required to explain why DA activity precedes BF bursting. Our data do not, however, exclude contributions from such inputs. Reward-prediction activity in both populations is likely shaped by distributed circuitry, and common inputs may still contribute to their coordinated recruitment^19,31–33^. Our findings therefore identify a causal route through which DA reward-prediction activity can rapidly influence BF bursting neurons, regardless of which upstream inputs shape this activity or how the reward-prediction signals are generated.

The matching millisecond timescales of natural DA-BF coupling and optogenetically evoked BF responses further link the experimentally demonstrated pathway to the coupling observed during behavior. Its persistence during acquisition, extinction, and contingency degradation suggests that DA-to-BF coupling is maintained as reward predictions are updated. The suppression of ongoing BF firing further suggests that DA-to-BF influence extends beyond the event-triggered propagation of phasic reward-prediction signals and provides ongoing excitatory drive to affected BF neurons.

Earlier studies documented dopaminergic projections from the VTA to the VP^20^ and rapid changes in VP activity following electrical stimulation of ventral tegmental regions^39^. Our findings extend this work by establishing that DA neurons exert rapid causal control over functionally identified BF bursting neurons. This strong functional influence is particularly striking given the comparatively sparse DA innervation of the substantia innominata and adjacent posterior VP relative to the striatum^22^.

DA manipulations nevertheless affected a substantial but incomplete subset of BF neurons, with BF bursting neurons enriched among the affected populations (Figures 4,5). The restriction of these effects to a subset of neurons could reflect limited manipulation efficacy due to incomplete viral expression or optical coverage, particularly at axon terminals. Alternatively, differences in DA innervation or dependence on DA input among BF bursting neurons could contribute to the variable effects. This possibility is consistent with the reported diversity of molecular identities, projection patterns, and functional properties among midbrain DA neurons^40–43^ and noncholinergic BF neurons^44–46^. Distinguishing these possibilities will require molecular markers that permit comprehensive identification and targeted manipulation of BF bursting neurons.

### A dopamine-receptor-independent mechanism for fast DA-to-BF coupling

A surprising mechanistic finding of this study is that DA-to-BF coupling occurs on a timescale of approximately 10 ms. This temporal relationship was conserved across rats and mice and across natural activity, optogenetic excitation, and optogenetic inhibition. Such rapid coupling is difficult to reconcile with canonical dopamine action through metabotropic D1 or D2 receptors, whose effects are mediated primarily through G-protein-dependent intracellular signaling cascades^23–25^. The persistence of fast BF recruitment during combined D1/D2 receptor blockade reveals a noncanonical mode of DA-neuron signaling that does not require activation of these receptors under the tested conditions. The shorter response latency following terminal relative to somatic stimulation further supports a local substrate within the BF rather than an indirect route through another DA target.

Glutamate cotransmission from DA neurons is the leading candidate mechanism. A subset of midbrain DA neurons expresses vesicular glutamate transporter 2 (VGLUT2) and other machinery required for vesicular glutamate release^35,47^, and fast glutamatergic transmission from DA terminals has been demonstrated across multiple forebrain targets^36,48–50^. In the BF, cotransmission would provide a parsimonious mechanism by which DA axons rapidly excite BF neurons while retaining the capacity for slower dopaminergic modulation. Consistent with this possibility, ATLAS tracing^38^ identified predominantly noncholinergic BF neurons as postsynaptic targets of VTA DA neurons at glutamatergic synapses. Establishing the causal contribution of glutamate cotransmission will require selective disruption of VGLUT2-dependent release from DA neurons combined with BF recordings.

Our findings suggest a distinct functional role for glutamate cotransmission from DA neurons. Previous studies have established fast glutamatergic transmission from DA terminals, particularly in the nucleus accumbens, but its reported behavioral effects have varied across studies and have sometimes been modest^35,37,48,51–53^. In the DA-to-BF pathway, glutamate cotransmission would enable DA neurons to recruit another subcortical broadcast system with millisecond precision, shaping both its phasic responses and ongoing activity. This would extend the functional role of DA glutamate cotransmission beyond local postsynaptic effects to coordinating the distribution of reward-prediction signals across brain systems.

### Coupled DA and BF outputs distribute reward-prediction signals to distinct downstream networks

DA-to-BF coupling provides a route through which DA reward-prediction activity can rapidly engage a second subcortical broadcast system with distinct downstream targets and functions. In the striatum, DA signaling contributes to synaptic plasticity, reinforcement learning, value updating, and action selection^13,15,54,55^. BF bursting neurons provide a separate route to cortical networks, where their activity has been linked to cortical activation, decision speed, and selective attention^7,8,16–18,26^. Coupling these populations may therefore allow the same predictive event to support both learning, through DA-dependent updates to values and actions, and current performance, through BF modulation of cortical processing.

More broadly, DA-to-BF coupling suggests that subcortical broadcast systems can coordinate their activity through rapid causal interactions. Such coupling may allow a shared reward-prediction signal to engage complementary functions across brain systems without requiring each system to compute that signal independently.

## Methods

### Animals and ethical approval

Experiments conducted at National Yang Ming Chiao Tung University (NYCU), Taiwan, were approved by the Institutional Animal Care and Use Committee (IACUC #1080510, #1131207, #1140404). Animal procedures conducted at Bordeaux University complied with European Directive 2010-63-EU and were approved by the Bordeaux University Animal Care and Use Committee (license authorization 51206). Animal numbers, sex, experimental allocation, and contributions to individual figures are summarized in Table S1.

### Rats

Ten adult male Long-Evans rats (National Laboratory Animal Center, Taiwan), aged 3–6 months and weighing 350–500 g at the start of experiments, were used. Rats were housed under a 12-h light/dark cycle and received 18 g of food daily, with water available ad libitum before training. During training and recording, water access was restricted to maintain 85–90% of their original body weight. Daily behavioral sessions lasted 60 min during the light phase and were followed by 15 min of water access.

### Mice

A total of 24 adult heterozygous DAT-IRES-Cre mice (21 male and 3 female), derived from B6.SJL*-Slc6a3*^tm1.1(cre)Bkmn^/J founders (The Jackson Laboratory, RRID:IMSR_JAX:006660)^56^ and bred in-house on a C57BL/6J background, were used for electrophysiological and pharmacological-validation experiments. Mice were 8–20 weeks old at the start of experiments and were housed under a 12-h light/dark cycle, with the dark phase from 07:00 to 19:00. Mice with electrode implants (n = 14; 12 male and 2 female) underwent daily behavioral sessions lasting 30–150 min during the dark phase. Water was available ad libitum except during behavioral training and recording, when mice received a total of 1 ml daily to maintain approximately 80% of their baseline body weight. This total included water obtained during behavioral sessions and supplemental water provided afterward^57^. The remaining mice (n = 10; 9 male and 1 female), used to assess the effects of dopamine receptor blockade under freely moving conditions, had unrestricted access to food and water.

The ATLAS tracing cohort comprised six additional female DAT-Cre mice, bred in-house on a C57BL/6J background. Mice were housed two to five per cage at 22–23°C and 40% relative humidity under a 12-h light/dark cycle, with food and water available ad libitum.

### Behavioral apparatus and tasks Rat behavioral apparatus

Behavioral tasks were conducted in custom-built operant chambers housed within light- and sound-attenuating cubicles, using the apparatus described previously^18^. Each chamber contained a central fixation port and two flanking reward ports; the right reward port was used for water delivery. Infrared sensors detected port entries and licking. Water was delivered through a solenoid-controlled sipper tube calibrated to dispense 10 μl per drop. Auditory stimuli were delivered through a speaker above the fixation port. A white LED above the speaker served as the intertrial-interval indicator in the lick-schedule reward task and as the novel visual cue in the learning experiments. Tasks were controlled by Med-PC software (Version V; Med

Associates, USA), which logged behavioral events at 1-ms resolution and sent TTL signals to the electrophysiological recording system for synchronization.

### Rat lick-schedule reward task

Rats first learned to obtain water by licking the right reward port, then to associate an auditory cue (80dB 6-kHz pure tone or 70dB white noise; counterbalanced across rats) with five 10-μl water drops delivered after the third lick within 5 s of cue onset. Cues were separated by unsignaled intertrial intervals (ITIs; 5–14 s), which were prolonged by licking.

Rats were subsequently trained on the fixation-based lick-schedule reward task (Figure 1B). A white LED illuminated the ITI (3–6 s); LED offset signaled the availability of a new trial. Fixation-port entries or licking during the ITI reset the interval. Rats initiated trials by entering the central fixation port and maintaining fixation for a pseudorandom foreperiod of 0.35, 0.50, 0.65, or 0.80 s. Premature fixation-port exit reset the trial.

Three trial types were presented pseudorandomly with equal probability: short-delay sound (S_SD_), long-delay sound (S_LD_), and catch trials. S_SD_ was the previously learned cue; S_LD_ was a distinct sound selected from a 6-kHz pure tone, white noise, or clicker, with cue identities counterbalanced across rats. Both sounds predicted three 10-μl water drops. Reward was delivered after the third lick on S_SD_ trials and after a session-specific threshold of 15–36 licks on S_LD_ trials. The latter threshold was progressively increased during training until rats behaviorally distinguished the two cues. Catch trials contained neither sound nor reward. Reaction time was defined as the interval from sound onset to fixation-port exit. Behavioral analyses were restricted to electrophysiological recording sessions only.

### Rat reward-learning experiments

A separate cohort completed the initial licking and auditory-association training described above, using a 6-kHz pure tone as the reward-predicting sound. Rats then underwent acquisition, extinction, and contingency degradation in sequence (Figure 3A). Throughout all phases, the familiar sound retained its reward contingency, providing an internal control and a consistent stimulus for functional identification of putative DA neurons and BF bursting neurons.

During acquisition (Experiment 1), a novel visual cue (white LED) predicted the same reward as the familiar sound. Sound and light trials were presented pseudorandomly with equal probability, separated by ITIs of 5–14 s. After performance stabilized, extinction (Experiment 2) began: reward was omitted on light trials, while stimulus probabilities and the sound–reward contingency remained unchanged. During contingency degradation (Experiment 3), the light was omitted. In its former trial slots, spontaneous licking could trigger an extra reward with 50% probability within an uncued 5-s response window. Sound trials continued to predict reward as before. For each learning phase, analyses used the session showing the largest within-session behavioral change in response probabilities.

### Mouse head-fixed behavioral apparatus

Behavioral experiments were conducted in a custom head-fixed setup housed within light- and sound-attenuating cubicles (55 × 43.5 × 54 cm, length × width × height). Each cubicle contained an exhaust fan that also masked external noise (Sanyo Denki America Inc., USA; Cat. No. 9S1224M401). Mice were secured by a metal head plate above a custom behavioral platform. Licking was detected by an infrared sensor positioned in front of the tongue (emitter: ams OSRAM, Germany; Cat. No.

SFH 4851; receiver: Vishay, USA; Cat. No. BPW76A). Water was delivered through a spout positioned in front of the mouse by a syringe pump calibrated to dispense 0.6 μl per motor step.

Auditory cues were delivered through a speaker positioned in front of the mouse (Visaton, Germany; Cat. No. 2240), controlled by a programmable audio generator (ANL-926; Med Associates, USA). A white LED lamp was positioned near the speaker (VCC, USA; Cat. No. VC1860245W3D). Med-PC software (Version V; Med Associates, USA) controlled the task, recorded behavioral timestamps at 1-ms resolution, and sent TTL signals to the electrophysiological recording and optogenetic stimulation systems.

### Mouse auditory cue-reward task

Following 1–2 weeks of postoperative recovery, mice underwent water restriction as described above and were habituated to head fixation for 1–2 days. During initial shaping, each lick triggered a 2.4-μl water reward if at least 1 s had elapsed since the preceding lick. Total water delivery was limited to 1 ml per session.

Once licking was stable, mice were trained to associate a 6- or 12-kHz pure tone (0.5 s, 80 dB), counterbalanced across mice, with this reward. It was delivered upon the third lick within a 3-s response window after cue onset. An unsignaled intertrial interval (ITI; 3–6 s) began after at least 1 s without licking; subsequent licking reset the ITI timer. The reward-predicting sound was used to identify BF bursting neurons during electrophysiological recordings.

For inhibition experiments, sound and catch trials were presented pseudorandomly with equal probability. Sound-reward was delivered upon the third lick within a 1.5-s response window after sound onset. Catch trials omitted the sound. Optogenetic inhibition was delivered on a randomly selected half of each trial type, allowing within-session comparisons with non-inhibited trials.

### Stereotactic surgery and viral injections

#### Rat electrode implantation

After initial behavioral training, rats underwent chronic electrode implantation using the electrode construction and general implantation procedures described previously^18^. Water restriction was suspended before surgery. Rats were anesthetized with isoflurane (0.5–2%; flow rate, 0.5–1.0 L/min) and received atropine (0.01 mg/kg, intramuscularly) to reduce respiratory secretions. Four skull screws anchored the implant: two cerebellar screws served as separate references for BF and VTA recordings, and two additional screws were placed over the frontal cortex. Electrical grounds were connected to the electrode cannulae.

Rats were implanted with custom microdrive assemblies for simultaneous BF and VTA recordings. Each electrode bundle contained 8–16 polyimide-insulated tungsten wires (38 μm diameter; California Fine Wire, USA) enclosed in a 28-gauge stainless-steel cannula and advanced using a precision microdrive. Two bundles targeted the BF bilaterally (AP, −0.5 mm; ML, ±2.25 mm; DV, −8.0 mm relative to bregma), with one bundle implanted in each hemisphere. VTA bundles targeted AP−5.4 mm, ML ±0.8 mm, and DV −8.2 mm relative to bregma. Three VTA implant configurations were used to accommodate unilateral or bilateral recordings and broaden recording coverage. One rat received a single vertical bundle targeting the unilateral VTA. Other rats received 10° angled bundles targeting either the bilateral VTA or a broader region of the unilateral VTA. In the latter configuration, two bundles within a shared microdrive were separated by 1 mm along the mediolateral or anteroposterior axis. Histological reconstruction confirmed that at least one bundle reached the VTA in every included rat.

Electrode impedances before implantation were approximately 0.1–0.2 MΩ, measured at 1 kHz using an Open Ephys Acquisition Board^58^. Cannulae were lowered at 2–50 μm/s using a robotic stereotactic manipulator (Neurostar GmbH, Germany) to the depth 1 mm above the target depth for BF or VTA separately. Implants were secured with dental acrylic (Hygenic Denture Resin, The Hygenic Corp., USA), and electrode tips were advanced a further 1 mm to spread the wires within the target regions. Rats received ibuprofen or acetaminophen for at least 3 days after surgery, with daily body-weight monitoring. Water restriction and behavioral training resumed 2 weeks after surgery.

### Mouse viral injections and implantation

DAT-Cre mice underwent surgery under isoflurane anesthesia (0.5–1.5%; flow rate, 0.5–1.0 L/min). Eyes were protected with ophthalmic ointment, the scalp was disinfected with 70% ethanol followed by 10% povidone-iodine, and the skull was leveled by aligning bregma and lambda.

For DA activation or inhibition, AAV-DIO-ChR2-EYFP (serotype 5 or 9; Addgene #20298) or AAV5-DIO-eNpHR3.0-EYFP (Addgene #26966), respectively, was injected bilaterally into the VTA (50–300 nl per site; titer, ≥1 × 10¹³ GC/ml). One female mouse received a 100-nl injection of a 1:1 mixture of AAV-DIO-ChR2-EYFP and AAV5-DIO-eNpHR3.0-EYFP^59,60^ and was included in both the DA activation and inhibition experiments. Injection sites were located at AP −3.1 mm and ML ±1.4 mm relative to bregma and 4.4 mm below the cortical surface, using a 12° mediolateral approach. Injections were delivered using a programmable nanoliter injector (Nanoject III; Drummond Scientific, USA) at 1 nl/s. The pipette was retained in place for 10 min before gradual withdrawal. For somatic manipulation, optic fibers were implanted using the same 12° mediolateral approach as the viral injections, targeting 4.05 mm below the cortical surface. A subset of mice also received VTA optrodes with adjustable optic fibers, constructed as described below, although only BF recordings were analyzed in this study. For manipulation of DA axon terminals in the BF, optic fibers were integrated into the BF optrodes described below.

Mice were implanted with bilateral BF optrodes consisting of two microdrive-controlled bundles, one in each hemisphere. Each bundle contained a 200-μm-core optic fiber (0.66 NA; Inper, China) surrounded by 14 polyimide-insulated tungsten wires (38-μm diameter; California Fine Wire, USA) and enclosed in 28-gauge polyimide tubing. Two optrode configurations were used. In most assemblies, the optic fiber remained fixed while the electrode wires could be advanced independently; during implantation, the fiber tip extended 1 mm beyond the retracted wire tips. In other assemblies, the optic fiber advanced together with the electrode wires, whose tips extended 250–500 μm beyond the fiber tip. Electrode impedances were approximately 0.1 MΩ at 1 kHz, measured using an Open Ephys Acquisition Board^58^.

BF electrodes targeted AP −0.1 mm and ML ±1.5 mm relative to bregma and 4.25 mm below the cortical surface. In most mice, the optrodes were implanted vertically. In a subset of mice, the cortical entry coordinates were adjusted and the optrodes were implanted using a 12° mediolateral approach to reach the same BF target region. During implantation, the electrode wires were retracted into the cannulae, which were lowered to 1 mm above the target depth using a micropositioner (Biochiefdom International Ltd., Taiwan), at 5 μm/s for the first 2 mm and 2 μm/s for the remaining 1.25 mm. The electrode wires were then advanced by 1 mm to reach the target depth.

Two pin connectors (#1372-2; Keystone Electronics, USA), positioned at AP −2.0 mm and ML ±3.0 mm relative to bregma, served as the electrical reference and ground. The electrode assemblies and skull screws were secured with Super-Bond (Sun Medical, Japan), and mice received a head bar for subsequent head-fixed experiments. Postoperative analgesia consisted of ibuprofen or acetaminophen for at least 3 days, with daily body-weight monitoring. Behavioral training began after 1–2 weeks of recovery, and optogenetic experiments began at least 4 weeks after surgery to allow viral expression.

### Electrophysiological recordings and spike sorting

Extracellular signals from the VTA and BF in rats and the BF in mice were recorded using an Open Ephys acquisition system^58^ at 30 kHz, with an acquisition bandwidth of 0.3 Hz–7.5 kHz. Signals were band-pass filtered at 250–6,000 Hz for offline spike sorting using KlustaKwik^61^, followed by manual curation in a custom Python-based graphical interface. Only single units with clusters clearly separated from noise, fewer than 0.1% of interspike intervals shorter than 1.5 ms were included. Cross-correlation analyses were used to identify and remove duplicate units recorded simultaneously across electrodes.

For rats, recording depths differed by at least 100 μm between included sessions, except during the reward-learning experiments. In these experiments, electrodes were advanced until stable recordings were obtained and then maintained at the same positions throughout each learning phase. For mice, recording depths differed by at least 62.5 μm between included sessions.

### Optogenetic activation and inhibition of VTA DA neurons and their BF terminals

For activation experiments, BF responses to optogenetic stimulation of VTA DA somata or their axon terminals in the BF were measured outside the task following the auditory cue-reward session. Blue light (465 nm; PlexBright; Plexon, USA) was delivered through implanted optic fibers at 3–6 mW, measured at the fiber tips. Stimulation consisted of either a single 10-ms pulse or 1-s trains of 10-ms pulses delivered at 10, 20, or 40 Hz. Light was delivered to the VTA for somatic activation and to the BF for terminal activation.

For inhibition experiments, light was delivered during a randomly selected half of sound and catch trials in the auditory cue-reward task. Illumination began at sound onset or the corresponding nominal onset on catch trials, allowing within-session comparisons with non-inhibited trials of the same type. Light from a 589-nm laser (MGL-F-589; CNI, China) or a 620nm LED (PlexBright; Plexon, USA) was delivered through implanted optic fibers at 4–5 mW, measured at the fiber tips. Inhibition lasted 200 ms in most sessions and 500 ms in a subset, with the same duration used for sound and catch trials within each session.

### Dopamine receptor blockade and locomotor assessment

D1-receptor antagonist SCH23390 hydrochloride (#HY-19545A, MedChemExpress, USA) and D2-receptor antagonist raclopride (#HY-103414, MedChemExpress, USA) were dissolved in PBS containing 1% DMSO and co-administered intraperitoneally at 0.4 and 2.0 mg/kg, respectively. Vehicle controls received PBS containing 1% DMSO, referred to as PBS in the figures. These doses were selected based on their established effects on striatal firing and LFP activity^34^. Head-fixed mice first performed the auditory cue-reward task, after which BF responses to somatic DA stimulation were measured before injection and at 10-min intervals for 60 min afterward using the stimulation protocols described above. Mice were briefly anesthetized with isoflurane during antagonist or vehicle injection to minimize distress while head-fixed. One male mouse used in DA somatic activation experiments under drug administration (D1/D2 or PBS) was excluded in LFP analyses, due to large 60Hz noise.

The behavioral efficacy of receptor blockade was assessed in a separate cohort of mice receiving the same antagonist combination (n = 5) or vehicle (n = 5). Locomotion was measured in custom-built Plexiglas chambers (23.8 × 18 × 25.7 cm; LabTalk Ltd., Taiwan) housed within light- and sound-attenuating cubicles with exhaust fans providing background noise. Mice moved freely without task demands or reward delivery for 30 min before injection and 60 min afterward. Position was tracked using DeepLabCut^62^, and the distance traveled in each 5-min bin was normalized to the mean distance across all pre-injection bins (Figure S9).

### Tissue preparation, histological verification, and reconstruction

For rats, animals were deeply anesthetized and transcardially perfused with normal saline followed by 10% formalin. Brains were postfixed with the implants attached for at least 24 h at 4°C, after which the implants were removed. Tissue was cryoprotected in 20–30% sucrose in PBS and sectioned coronally at 50 µm using an SM2010 R microtome (Leica Biosystems, Germany). Electrode-tip locations were verified by dark-field microscopy using an M205 FCA microscope (Leica Microsystems, Germany).

For mice with electrode implants, animals were deeply anesthetized and transcardially perfused with PBS followed by 4% paraformaldehyde. Brains were cryoprotected and sectioned as described for rats. To visualize DA neurons and augment visualization of viral expression coverage, some sections were immunostained for tyrosine hydroxylase (TH) and GFP . After PBS wash, sections were first incubated in PBS containing 0.2% Triton X-100 (PBST) for 40-60 min, and blocked for 2 h at room temperature in PBS containing 3% normal donkey serum (Jackson ImmunoResearch, USA; RRID: AB_2337258). Next, sections were incubated with primary antibodies, rabbit anti-TH (1:200; cat. no. AB152; Sigma-Aldrich, USA) and chicken anti-GFP (1:1000; cat. no. ab13970; abcam, UK) for 12-48 h at 4°C. After washing, sections were incubated with secondary antibodies, donkey anti-chicken Alexa Fluor 488 (1:1000; cat. no. A-78948; RRID: AB_2921070; Invitrogen, USA) and goat anti-rabbit Alexa Fluor 647 (1:500; cat. no. A-21245; RRID: AB_2535813; Invitrogen, USA), for 2 h at room temperature. After washing, sections were mounted with VECTASHIELD Antifade Mounting Medium containing DAPI (Vector Laboratories, USA) and examined by fluorescence microscopy (Carl Zeiss, Germany) to verify viral expression and implant locations.

Rat histological sections were registered using QuickNII^63^ to version 4.01 of the Waxholm Space Sprague Dawley rat brain atlas^64,65^. Mouse recording sites, implant tracks, and viral expression were reconstructed with reference to the Allen Mouse Brain Common Coordinate Framework v3.0^66^.

### ATLAS transsynaptic tracing and cell-type quantification

To label BF neurons postsynaptic to VTA DA neurons through glutamatergic synapses, DAT-Cre mice received unilateral injections of AAV5-hSyn-DIO-ATLASsnFLP (120 nL; Addgene #232352; produced by the IMN Vector Core) into the VTA and AAV8-EF1α-fDIO-mCherry (120 nL; Addgene #114471-AAV8) into the ipsilateral BF. VTA coordinates were AP −3.20 mm and ML−0.50 mm relative to bregma and DV −4.20 mm from the brain surface. BF coordinates were AP +0.10 mm, ML −1.50 mm, and DV −4.65 mm. Four weeks after injection, mice were transcardially perfused with PBS. Brains were immersion-fixed in 4% paraformaldehyde for 48 h and stored in PBS containing 0.03% sodium azide.

Coronal sections (40 µm) were cut on a vibratome (Leica Biosystems, Germany), washed in PBS, and blocked for 90 min at room temperature in 10% normal donkey serum prepared in PBS containing 0.3% Triton X-100 (PBST). Sections were incubated for 72 h at 4°C with goat anti-ChAT (1:200; cat. no. AB144P; RRID: AB_2079751; Sigma-Aldrich, USA) and rat monoclonal anti-RFP (1:1,000; clone 5F8; cat. no. 5f8; RRID: AB_2336064; ChromoTek, Germany) primary antibodies. After washing, sections were incubated for 2 h at room temperature with donkey anti-goat IgG conjugated to Alexa Fluor 488 (1:500; cat. no. A-11055; RRID: AB_2534102; Invitrogen, USA) and donkey anti-rat IgG conjugated to Alexa Fluor 594 (1:500; cat. no. A-21209; RRID: AB_2535795; Invitrogen, USA) secondary antibodies. Nuclei were labeled with Hoechst 33342 (1:10,000; cat. no. H3570; Invitrogen, USA) for 5 min.

Images were acquired using an SP5-MP confocal microscope (Leica Microsystems, Germany). For each mouse (n = 6), one image was acquired at 20× from each of four AP levels of the right BF. RFP-positive neurons were counted manually and classified as ChAT-positive or ChAT-negative using the Cell Counter plugin in Fiji.

### Data analysis

Data were analyzed using custom MATLAB scripts (R2020a; The MathWorks Inc., USA).

### Functional identification of putative DA neurons and BF bursting neurons

Putative DA neurons in rats and BF bursting neurons in rats and mice were identified using shared functional criteria: a baseline firing rate 0.1–10 spikes/s and a phasic response to a reward-predicting sound. PSTHs were constructed using 10-ms bins, and sound responsiveness was defined as a mean firing rate during the [0.05, 0.2]-s post-cue window exceeding baseline by more than 2 spikes/s. These criteria were consistent with previous descriptions of the two populations^1,7,9,18,67^.

In rats, identification used responses to the short-delay sound (S_SD_) in the lick-schedule reward task or to the consistent, reward-predicting sound maintained throughout the reward-learning experiments. Baseline firing rates were measured from [-2, -1]-s interval relative to the trial-start signal in the lick-schedule reward task and 1 s prior to sound onset in the reward-learning experiments.

In mice, BF bursting neurons were identified by their responses to the reward-predicting sound in the auditory cue-reward task. Baseline firing rates were measured during the 1 s preceding sound onset.

### Hierarchical clustering of neuronal response profiles

To assess whether the functional identification criteria corresponded to coherent response-defined populations^67,68^, we performed hierarchical clustering separately for all VTA and BF neurons recorded during the rat lick-schedule reward task (Figure S2). For each neuron, baseline-subtracted PSTHs were constructed using 10-ms bins from −0.3 to 0.3 s around trial start, short-delay sound (S_SD_) onset, and the corresponding nominal sound onset on catch trials. The three PSTHs were concatenated into a single response vector.

Pairwise response similarity, r, was defined as the maximum cross-correlation coefficient between response vectors at lags within ±0.1 s. Allowing these small temporal offsets accommodated variation in response latency across animals and sessions. A dissimilarity matrix defined as 1−r was subjected to agglomerative hierarchical clustering using average linkage, following Avila and Lin^68^. For visualization, dendrogram branches were reordered according to each neuron’s mean squared similarity to all neurons in the population without altering the tree topology. Functionally identified putative DA neurons and BF bursting neurons were marked alongside the similarity matrices and ordered response heatmaps to assess their correspondence with the resulting clustering organization.

### Trial-by-trial response amplitude correlations

For each neuron, response amplitude was quantified as the mean firing rate within the following event-aligned windows: trial-start signal, 80–250 ms; visual stimulus onset, 80–250 ms; auditory stimulus onset, 50–200 ms; and reward delivery, 50–200 ms.

For population correlations (Figures 1I and 3D), response amplitudes were averaged across simultaneously recorded neurons within each population on each trial. Within each session, Pearson’s r was calculated across trials for four population pairings: putative DA neurons with BF bursting neurons, putative DA neurons with other BF neurons, other VTA neurons with BF bursting neurons, and other VTA neurons with other BF neurons. Within-session correlations were considered significant at p < 0.01. For each event or epoch, session-wise correlation coefficients were tested against zero using two-sided Wilcoxon signed-rank tests and compared between population pairings using one-sided Wilcoxon signed-rank tests. P values were Bonferroni-corrected within each family of comparisons.

For individual-neuron-pair correlations (Figure S3), Pearson’s r was calculated across trials between the response amplitudes of each simultaneously recorded VTA–BF neuron pair. Pairs were organized into the same four categories used for population analyses. Significant positive and negative correlations were defined at p < 0.01.

### Spike-time cross-correlation analysis

Spike-time cross-correlograms were computed for all simultaneously recorded VTA-BF neuron pairs using 1-ms bins over lags from −3 to 3 s. VTA spikes served as the reference, with positive lags indicating VTA-leading activity. Each cross-correlogram was normalized as (observed − expected)/expected. This measure expresses the relative excess in coincidence counts, rather than a Pearson correlation coefficient: zero indicates the expected count, and a value of 1 indicates twice that count.

For analyses using all session spikes, the expected coincidence count was estimated as the mean count across baseline lags from −3 to −1 s and 1 to 3 s. For ITI analyses, expected counts at each lag were estimated by averaging 1,000 circularly shuffled cross-correlograms. Shuffling introduced a random temporal offset between the spike trains while preserving the ITI epoch structure .

A neuron pair was classified as significantly correlated if at least 10 bins within ±0.3 s fell outside the range bounded by the minimum and maximum cross-correlation values at baseline lags^16^. Cross-correlograms were then smoothed using a 10-bin

Hanning window for visualization, and peak lags were estimated from the smoothed cross-correlograms of significant pairs. Significant pairs were used to calculate the population-average cross-correlograms and peak-lag distributions in Figures 2A,B and 3E. For the analyses in Figures 2A,B and the quantification of significant-pair proportions in Figure S4, only rats contributing at least five putative DA neuron–BF bursting neuron pairs were included (n = 5 rats). Pairs identified as significant using all spikes recorded during the session in Figure 2A were also used for the population noise-correlation analysis in Figure 2D.

### Noise correlation analysis

Joint peri-stimulus time histogram (jPSTH) analysis was performed using the MATLAB jpsth toolbox following established procedures^69,70^. Spike counts were computed in 10-ms bins from −0.5 to 0.5 s around event onset, separately for each trial condition.

For each simultaneously recorded VTA–BF neuron pair, the raw jPSTH contained the mean number of within-trial VTA–BF spike pairs for every combination of VTA and BF time bins. The shift predictor was calculated as the outer product of the neurons’ mean PSTHs and represented the expected spike-pair count attributable to their shared event-locked responses. The shift predictor was subtracted from the raw jPSTH, and each resulting element was divided by the product of the across-trial spike-count standard deviations at the corresponding time bins, yielding the normalized jPSTH, or noise correlation matrix.

For each pair, averaging along matrix diagonals corresponding to constant time differences produced a noise correlation lag profile, with positive lags indicating VTA-leading activity. Profiles were summarized as mean ± s.e.m. across pairs within each category in Figure 2E.

### Directly excited BF neurons: identification and response characterization

To identify BF neurons directly excited by optogenetic activation of DA neurons, PSTHs were constructed using 1-ms bins aligned to the first stimulation pulse. Responses to the first pulse were pooled across stimulation protocols; subsequent pulses within stimulation trains were not used for classification. A neuron was classified as directly excited if at least one bin within 0–25 ms after stimulation onset exceeded the maximum PSTH bin value observed during the −1 to 0-s prestimulation baseline. This criterion was applied to both the DA somatic- and terminal-stimulation experiments.

For directly excited BF neurons, evoked spiking was further characterized by onset latency and spiking probability following each stimulation pulse. A pulse was considered to evoke a spike if at least one spike occurred within 25 ms after pulse onset. For pulses that evoked a spike, onset latency was defined as the interval from pulse onset to the first spike. Spiking probability was calculated as the proportion of stimulation trials in which a spike occurred within this window.

### Directly inhibited BF neurons: identification and latency estimation

Directly inhibited BF neurons were identified from stimulated catch trials, which lacked cue-evoked activity. PSTHs were constructed using 1-ms bins from −1 to 2 s relative to stimulation onset. Candidate neurons were first identified using a cumulative frequency histogram method^71^. Cumulative responses after stimulation onset were compared with null distributions generated from 1,000 bootstrap resamples of the −1 to 0-s baseline PSTH. Neurons were retained as candidates if their first significant modulation was inhibitory (α = 0.001) and exceeded the minimum absolute cumulative-response magnitude of 5.

A descending sigmoid was then fitted to the unsmoothed catch-trial PSTH of each candidate neuron from −0.5 to 0.2 s relative to stimulation onset. Because sparse firing and small absolute changes in firing rate can delay significance-based estimates of inhibition onset, the sigmoid transition point was used to estimate latency. Neurons with transition points within 25 ms after stimulation onset were classified as directly inhibited. The same criteria were applied to inhibition of VTA DA somata and their axon terminals in the BF.

To estimate population inhibition onset, unsmoothed PSTHs were pooled across directly inhibited neurons for each stimulation site and fitted with the same descending sigmoid. The fitted transition point was taken as the population latency reported in Figure 5. The population traces in Figures 5G and 5M were subsequently smoothed using a 5-bin Hanning window for visualization only.

### BF LFP spectral analysis

BF LFP signals were downsampled from 30 kHz to 1 kHz and preprocessed to remove large-amplitude artifacts, 60-Hz line noise, and its harmonics. Channels containing artifacts (∣Z∣>10) during more than 20% of the recording were excluded. Power spectral density was calculated using 2-s windows with 1-s overlap and a frequency resolution of 0.5 Hz, then averaged across included BF channels within each session. Power at each frequency was normalized to its mean during the baseline period, defined as all valid recording periods before the 1-min interval immediately preceding the onset of isoflurane anesthesia, with optogenetic-stimulation epochs excluded.

For each time bin, frequency-normalized power was averaged within the low-frequency (4–15 Hz) and high-frequency (30–60 Hz) bands. The time-resolved power ratio was calculated by dividing the high-frequency band average by the low-frequency band average (Figure S9A,B). Power ratios were then averaged over the baseline and 5–60-min post-injection periods for session-level comparisons (Figure S9C).

## Statistical analysis

Pairwise comparisons were corrected for multiple comparisons using the Bonferroni method. Adjusted p values are reported.

---

**Figure S1.**
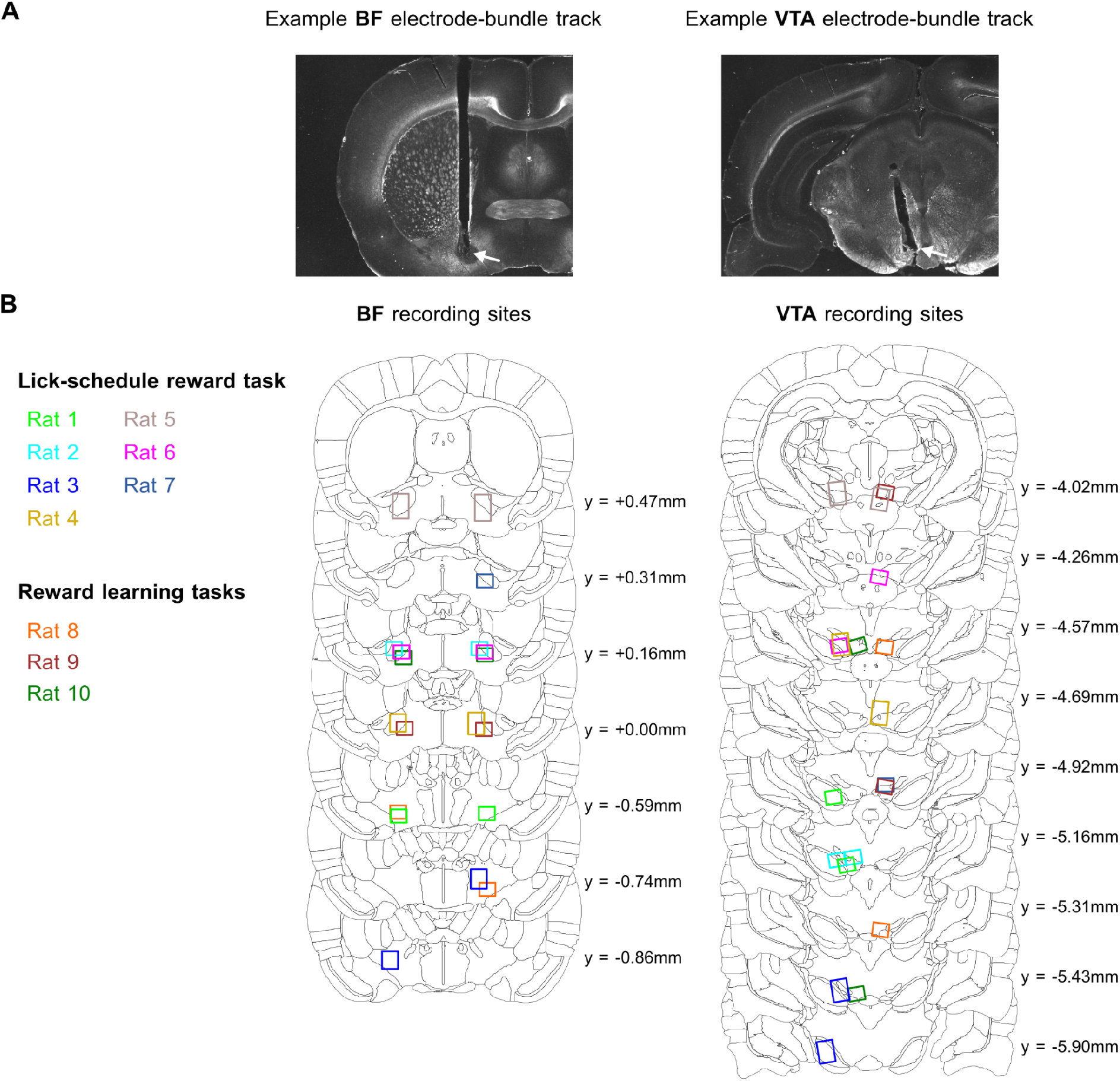
Histological verification of BF and VTA recording sites in rats. **A**, Representative dark-field images of coronal sections showing tracks of electrode bundles in the BF (left) and VTA (right). White arrows indicate bundle-tip locations. **B**, Histological reconstructions of BF (left) and VTA (right) recording sites on coronal sections of the Waxholm Space (WHS) rat brain atlas^64,65^. Colors identify individual rats from the lick-schedule reward task and reward-learning experiments. Each box represents the estimated location of an electrode bundle across the recording sessions included in the analysis. Labels indicate anteroposterior coordinates in WHS space. The number of VTA recording sites varied across rats because different electrode configurations targeted one or both hemispheres, with one or two bundles in a given hemisphere. Histological reconstruction confirmed that at least one bundle reached the VTA in every included rat.

**Figure S2.**
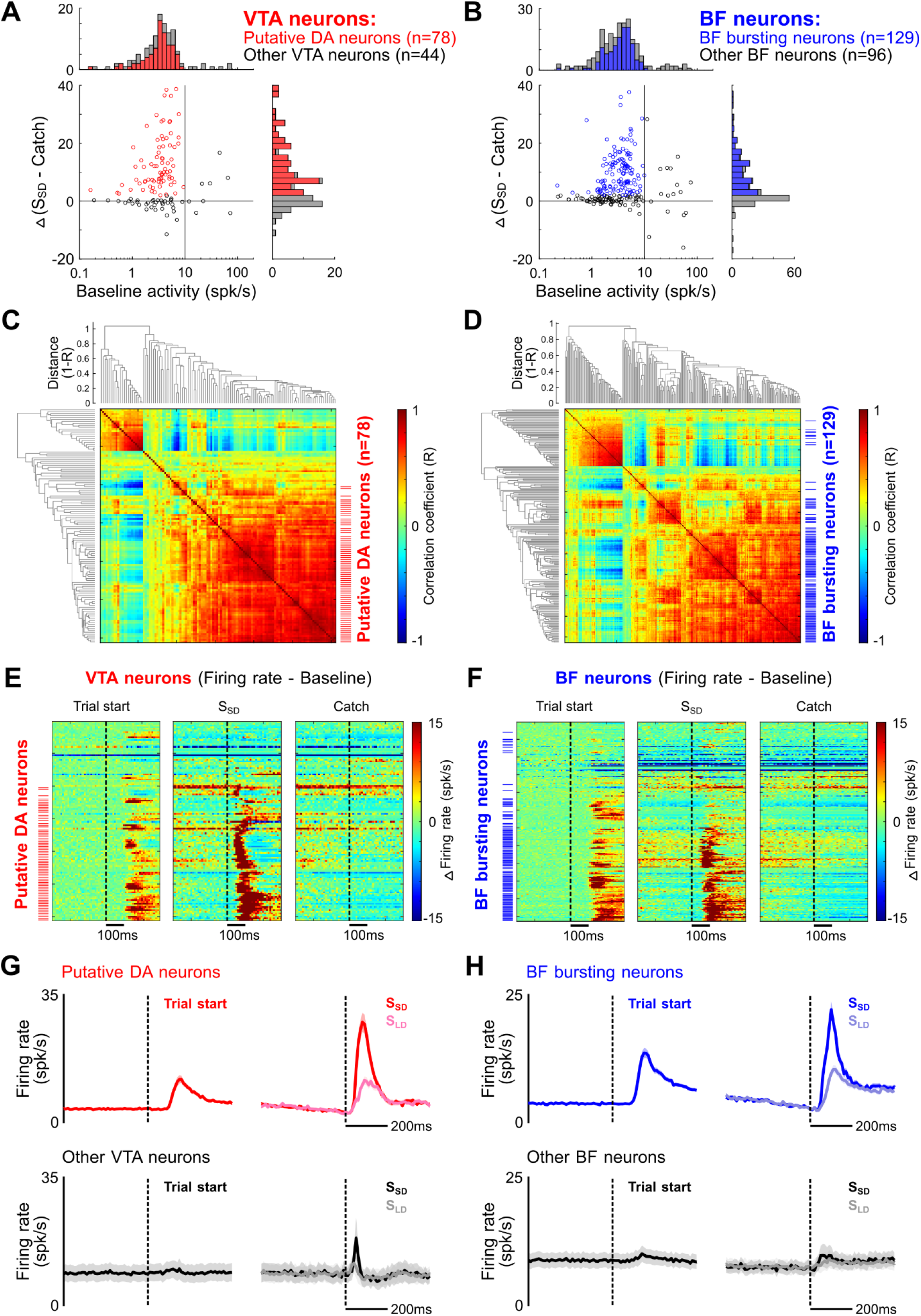
Identification of putative DA neurons and BF bursting neurons in the lick-schedule reward task. **A**, Operational identification of putative DA neurons using two established functional criteria: a baseline firing rate ≤10 spikes/s and a phasic response to the reward-predicting S_SD_ that exceeded the corresponding catch-trial response by at least 2 spikes/s within the burst window. The scatterplot shows both criteria for all VTA neurons, with marginal distributions. Red indicates putative DA neurons (n = 78), and gray indicates other VTA neurons. **B**, Corresponding identification of BF bursting neurons using the same criteria. Blue indicates BF bursting neurons (n = 129), and gray indicates other BF neurons. **C, D**, Hierarchical clustering of all VTA (**C**) and BF (**D**) neurons based on the similarity of their baseline-subtracted firing patterns aligned to trial start, S_SD_ onset, and the corresponding time in catch trials. Dendrograms show correlation distance (1 − R), and matrices show pairwise response correlations. Red and blue ticks indicate putative DA neurons and BF bursting neurons, respectively, as defined by the functional criteria in **A** and **B**. **E, F**, Baseline-subtracted responses of the same VTA (**E**) and BF (**F**) neurons, arranged in hierarchical clustering order. Each row represents one neuron, with the same neuronal order across trial start, S_SD_, and catch conditions. Dashed lines indicate event onset or the corresponding time in catch trials. Red and blue ticks indicate putative DA neurons and BF bursting neurons, respectively. **G, H**, Average PSTHs of putative DA neurons and other VTA neurons (**G**) and of BF bursting neurons and other BF neurons (**H**), aligned to trial start and the reward-predicting sounds S_SD_ and S_LD_. Traces show mean ± s.e.m. across neurons. Putative DA neurons and BF bursting neurons exhibited prominent phasic responses to trial start and reward-predicting sounds, with larger sound-evoked responses to S_SD_ than S_LD_. Other VTA and BF neurons showed substantially weaker event-related modulation. Together, these analyses show that the simple functional criteria capture coherent response-defined subsets and support their use as a common operational definition in subsequent experiments.

**Figure S3.**
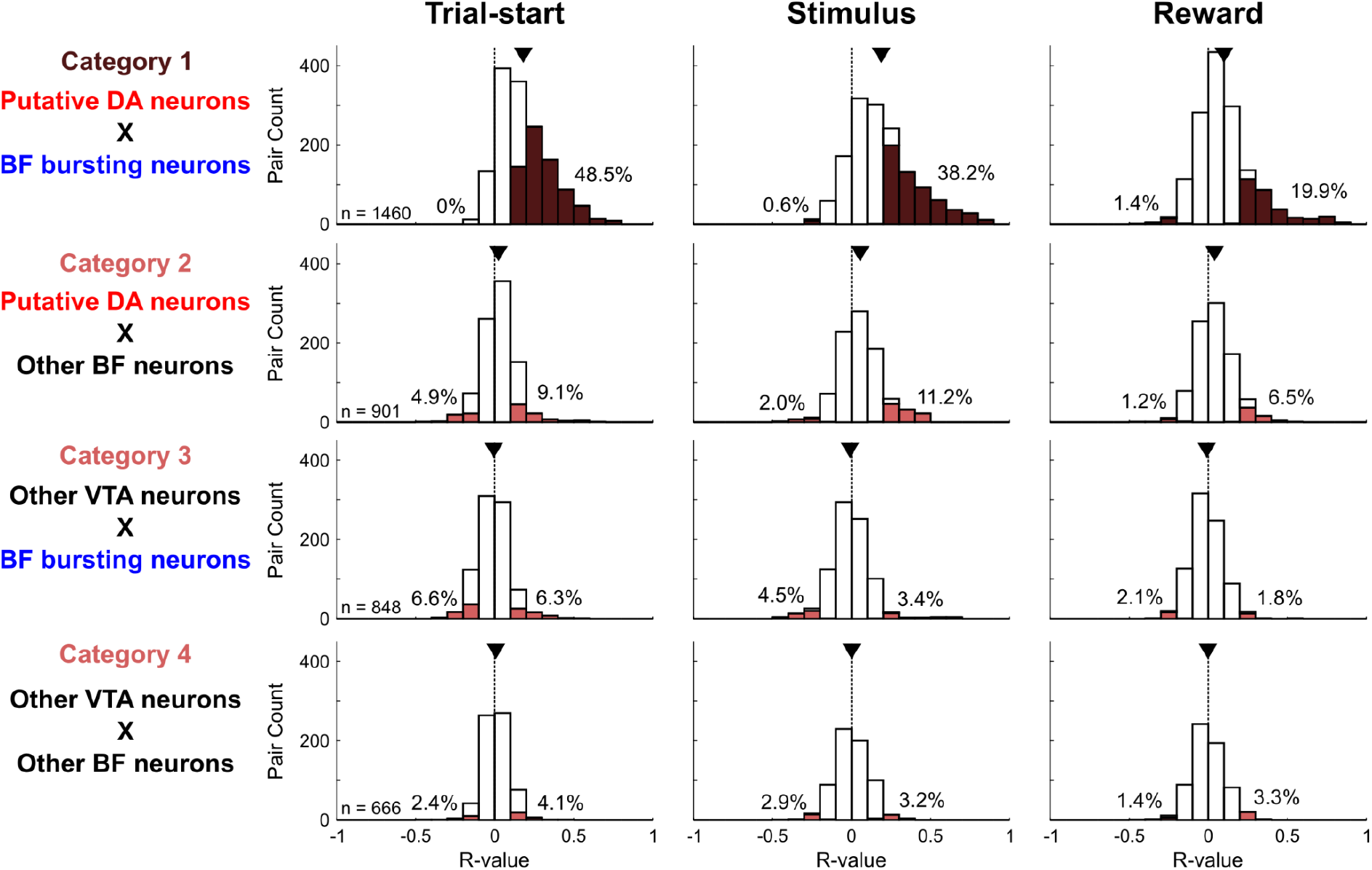
Trial-by-trial response-amplitude correlations are selective for putative DA neuron–BF bursting neuron pairs. Distributions of Pearson’s correlation coefficients for trial-by-trial response amplitudes between simultaneously recorded VTA and BF neuron pairs at trial start (left), reward-predicting stimulus onset (middle), and reward delivery (right). Rows show the four pairing categories defined in Figure 1I and used in Figure 3D. Each coefficient was calculated across trials for one neuron pair and event. Colored histogram portions indicate significant correlations (p < 0.01), and black triangles indicate distribution means. Percentages indicate the proportions of significant negative (left) and positive (right) correlations. Putative DA neuron-BF bursting neuron pairs showed a marked shift toward positive correlations across all three event types.

**Figure S4.**
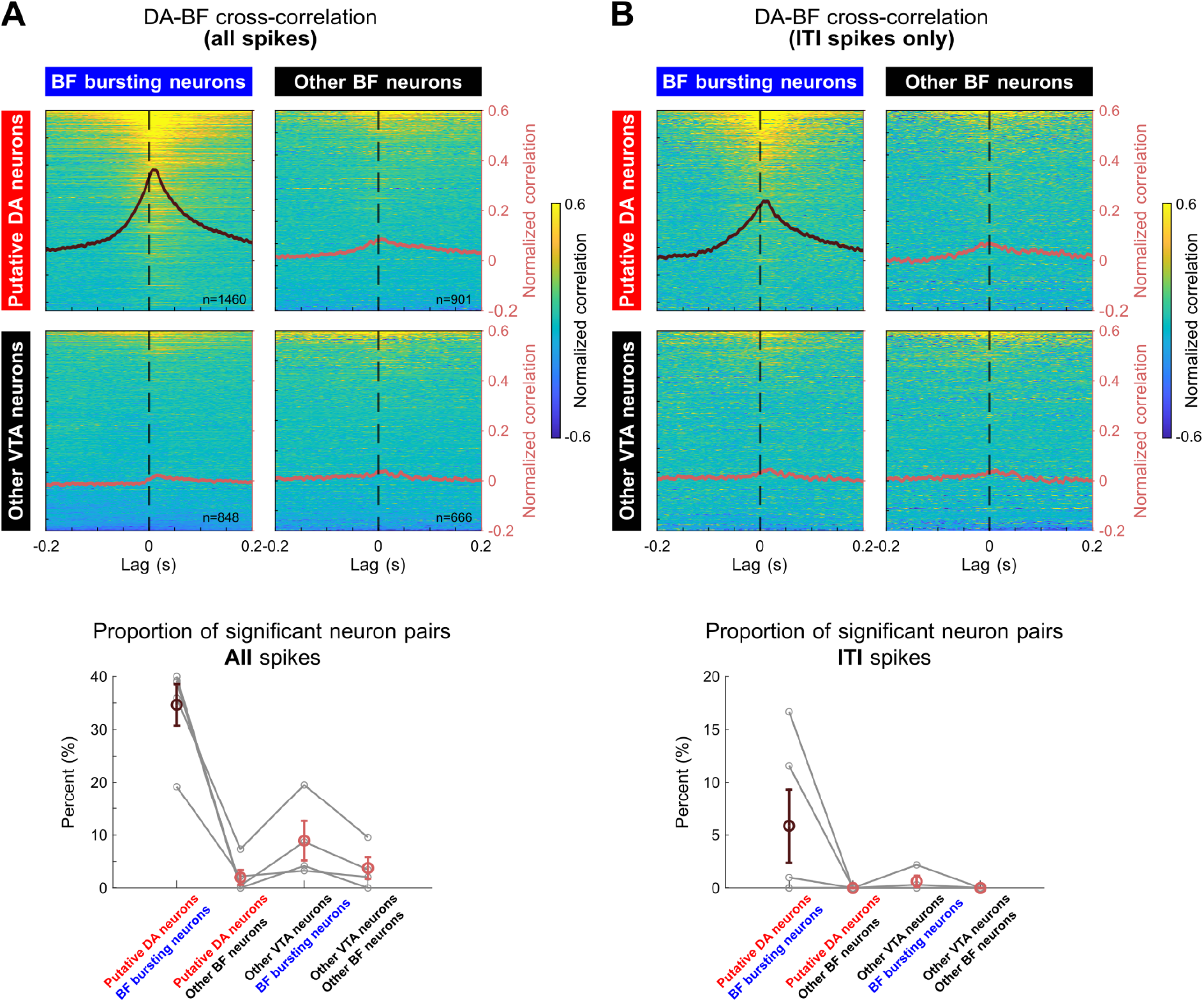
Selective spike-time coupling between putative DA neurons and BF bursting neurons. **A**, Normalized spike-time cross-correlations for simultaneously recorded VTA-BF neuron pairs, calculated using all spikes from each session. Cross-correlograms were normalized as (observed − expected)/expected, expressing the relative excess in coincidence counts rather than a Pearson correlation coefficient. Zero indicates the expected count, and a value of 1 indicates twice that count. Heatmaps show individual neuron pairs organized into the four population pairings defined in Figure 1I and sorted by mean correlation within ±0.3 s. Superimposed traces show the mean for each pairing. Bottom, percentages of significantly correlated pairs, as defined in Methods. Gray symbols and lines indicate individual rats (n = 5); colored symbols show mean ± s.e.m. **B**, Corresponding analyses restricted to ITI spikes. Significant putative DA neuron-BF bursting neuron pairs from **A** and **B** were used for the corresponding analyses in Figure 2A,B; pairs identified using all session spikes in **A** were also used in Figure 2D. Putative DA neuron-BF bursting neuron pairs showed the strongest cross-correlation peak and the highest proportion of significant pairs in both analyses, demonstrating selective temporal coupling outside overt task events. Lag conventions are as in Figure 2A.

**Figure S5.**
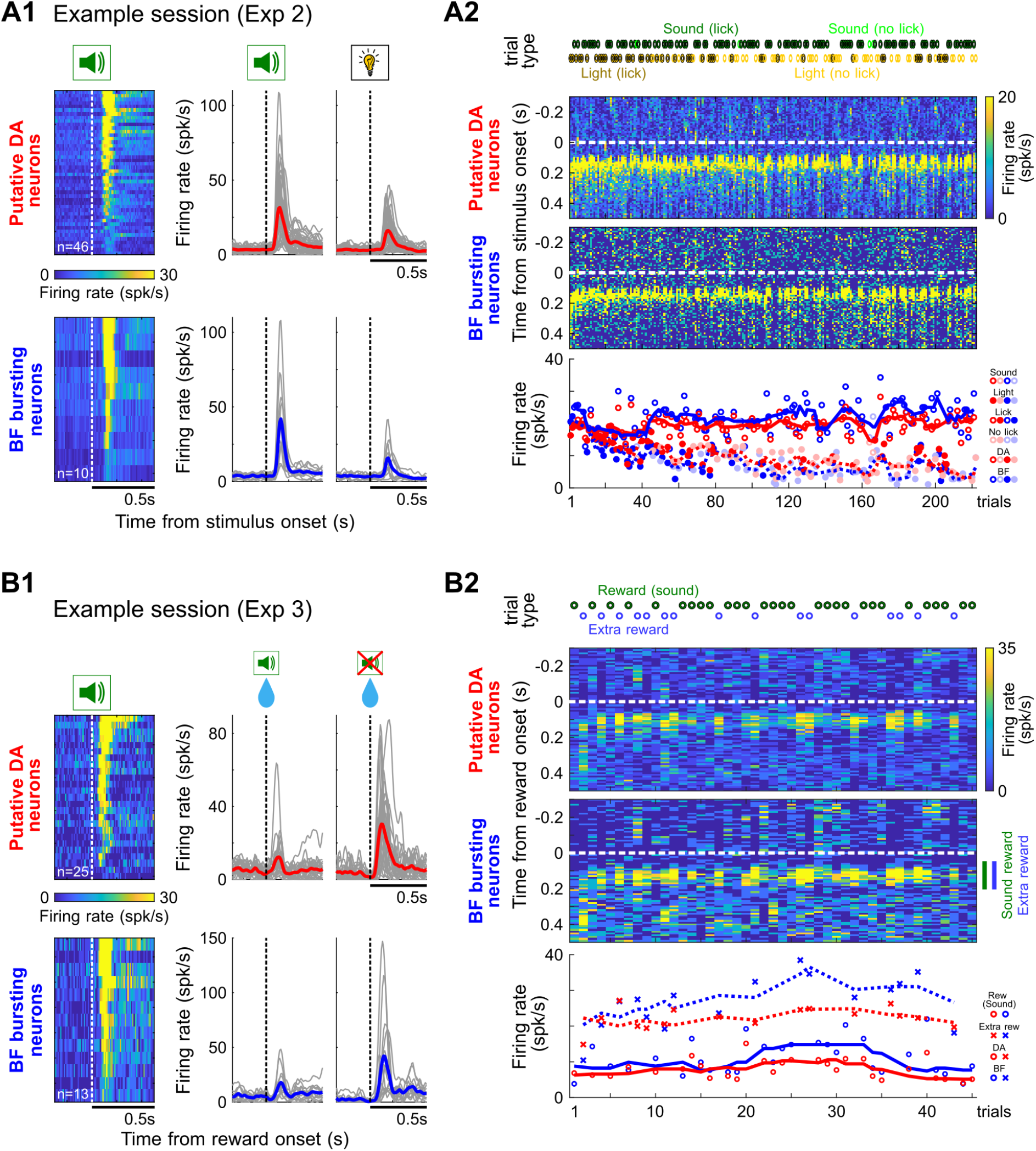
Coordinated putative DA and BF bursting population dynamics during extinction and contingency degradation. **A**, Evolution of putative DA and BF bursting activity during an example extinction session from Experiment 2. Conventions are as in Figure 3C. **A1**, Heatmaps show responses to the consistent sound used to identify putative DA neurons and BF bursting neurons. PSTHs show responses aligned to the consistent sound and extinguished light. **A2**, Trial-by-trial stimulus-aligned population activity and response amplitudes from the same session. DA and BF activity remained closely matched as responses to the extinguished light declined. **B**, Corresponding analyses **for** an example contingency-degradation session from Experiment 3. **B1**, Heatmaps show responses to the consistent sound used for functional identification. PSTHs show responses aligned to expected rewards on sound trials and unexpected extra rewards. **B2**, Trial-by-trial reward-aligned population activity and response amplitudes for expected and unexpected rewards. Colored bars indicate the response windows used for amplitude measurements. DA and BF activity remained closely matched across reward conditions, and both populations showed consistently larger responses to unexpected rewards than to expected rewards on sound trials.

**Figure S6.**
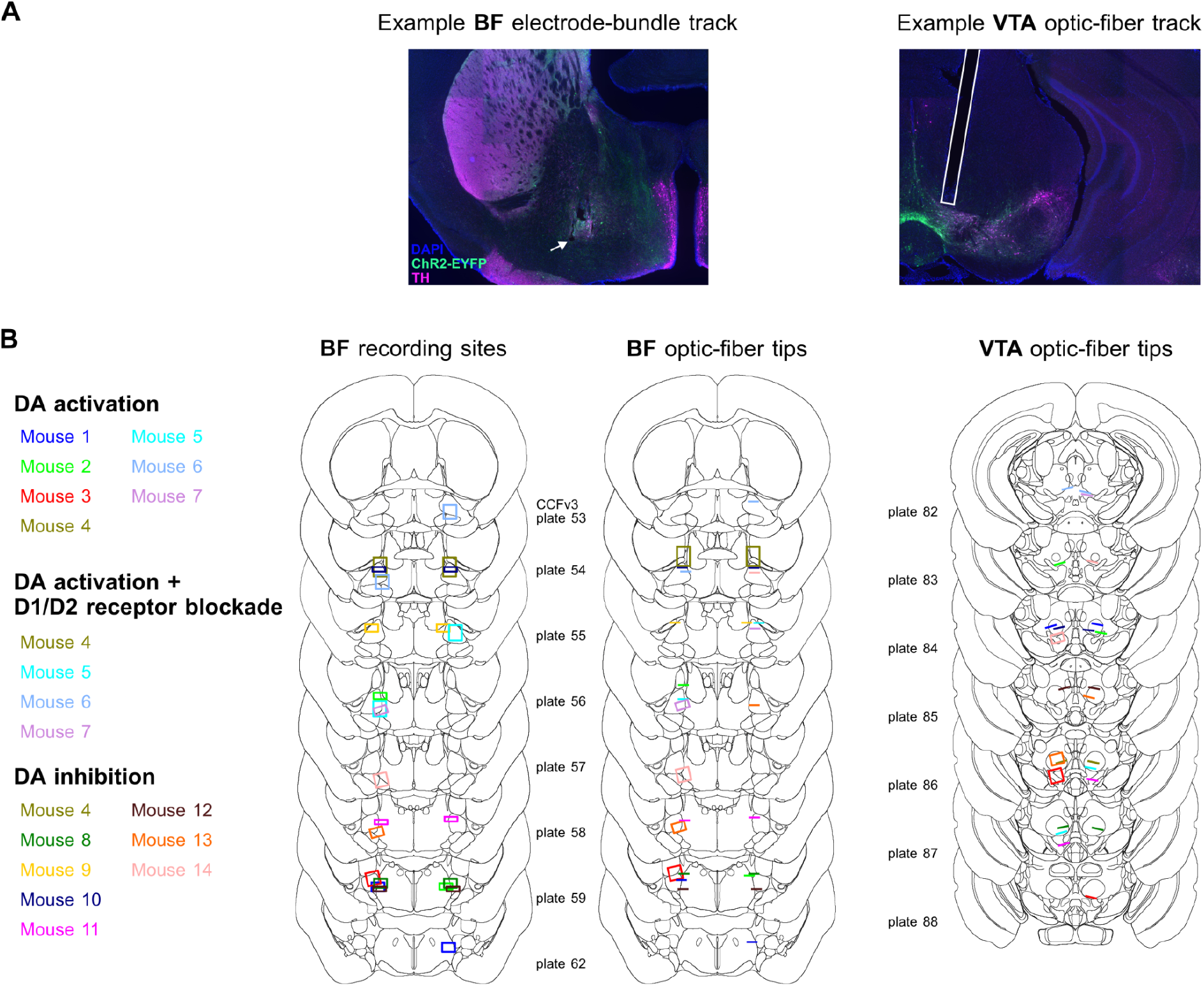
Histological verification of BF recording sites and optic-fiber placements in mice. **A**, Representative images showing an electrode-bundle track in the BF (left) and an optic-fiber track in the VTA (right). The white arrow indicates the electrode-bundle tip; the white outline marks the optic-fiber track. TH, tyrosine hydroxylase. **B**, Histological reconstructions of BF recording sites (left), BF optic-fiber tips (middle), and VTA optic-fiber tips (right) on coronal sections of the Allen Mouse Brain Common Coordinate Framework v3 (CCFv3)^66^. Colors identify individual mice, with experimental assignments listed at left. For recording sites, each box represents the estimated location of an electrode bundle across the recording sessions included in the analysis. For optic fibers, line segments indicate fixed tip positions, whereas boxes indicate the range of tip positions across the sessions included in the analysis for adjustable fibers. Labels indicate CCFv3 reference atlas plate numbers.

**Figure S7.**
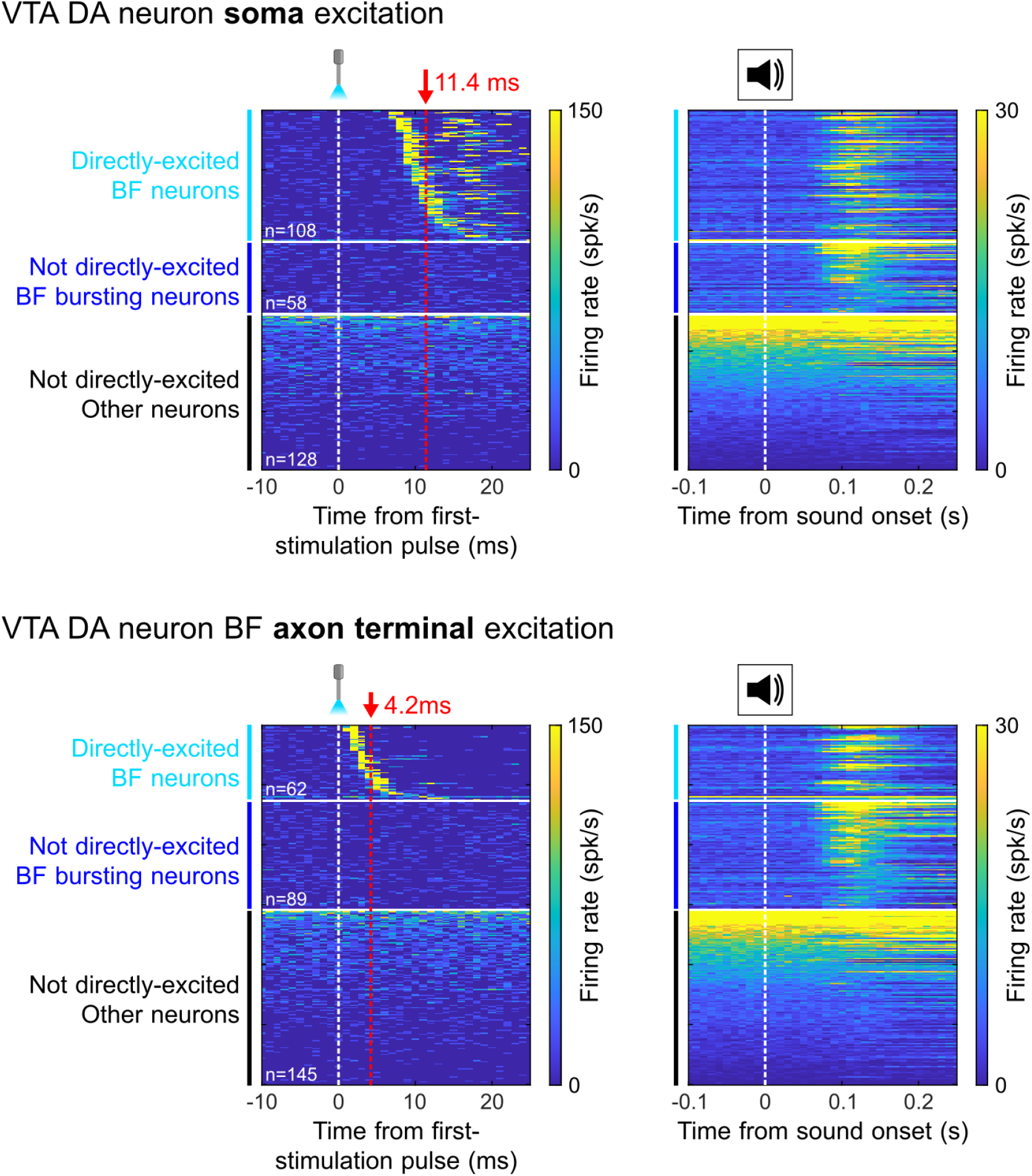
Population-wide BF response profiles during activation of VTA DA somata and their terminals in the BF. Heatmaps of all BF neurons recorded during activation of VTA DA somata (top) or their axon terminals in the BF (bottom). Responses to the first 10-ms light pulse (left) and the reward-predicting sound (right) are shown in the same neuronal order. Neurons were grouped as directly excited BF neurons, BF bursting neurons not directly excited, or other BF neurons not directly excited. Within these groups, neurons were sorted by first-spike latency, sound-response amplitude, and baseline firing rate, respectively. Red dashed lines indicate the mean first-spike latencies of directly excited neurons following somatic (11.4 ms) and terminal (4.2 ms) activation. Each row represents one neuron.

**Figure S8.**
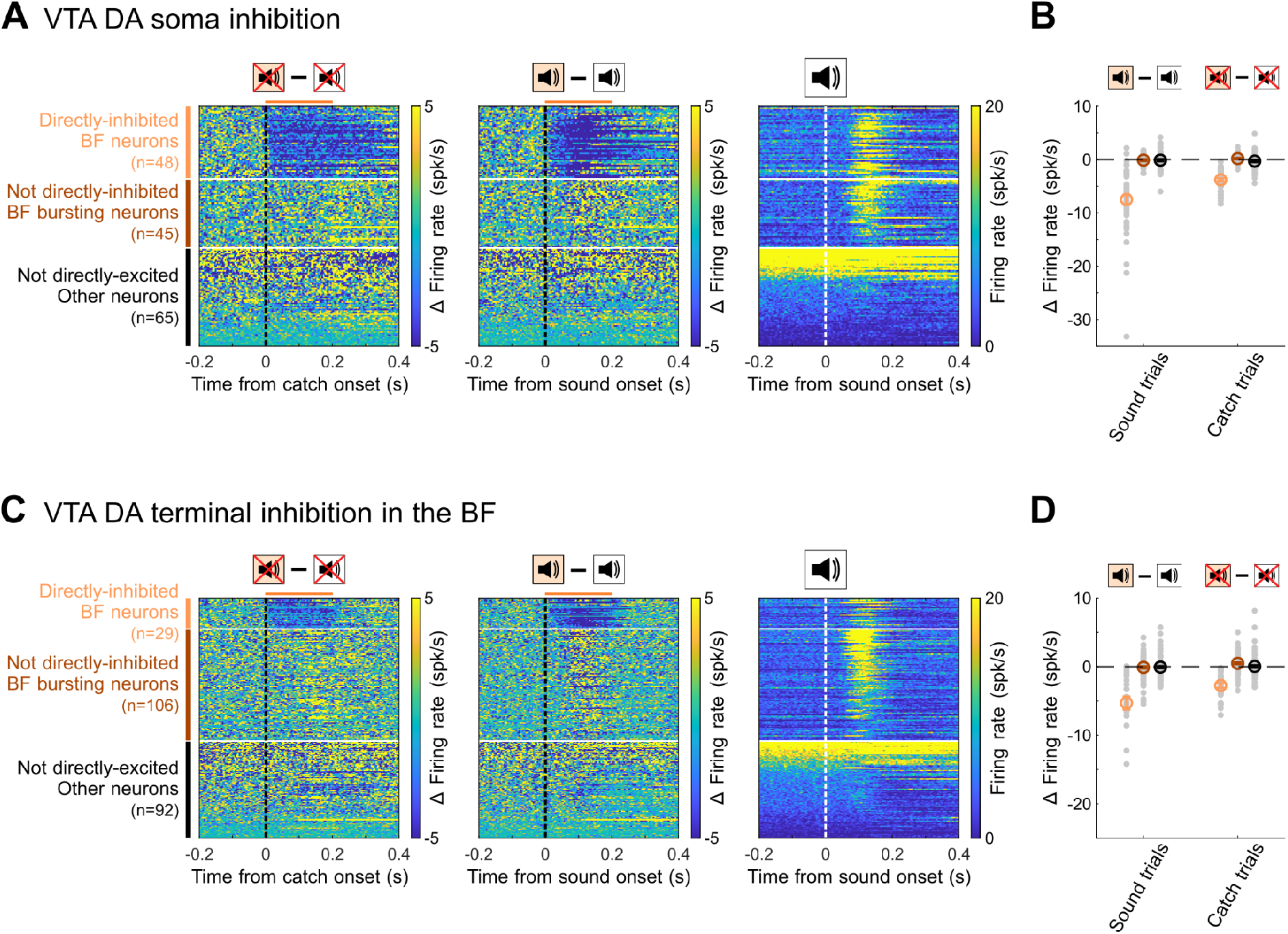
Population-wide BF response profiles during inhibition of VTA DA somata and their terminals in the BF. **A**, BF population responses during somatic inhibition of VTA DA neurons. Heatmaps show firing-rate differences between inhibited and non-inhibited trials (inhibited minus non-inhibited) on catch trials (left) and sound trials (middle), together with sound-evoked firing rates used to identify BF bursting neurons (right). Horizontal bars mark the 200-ms inhibition period common to all sessions; inhibition lasted 500 ms in a subset of sessions on both catch and sound trials. Neurons were grouped as directly inhibited BF neurons, BF bursting neurons not directly inhibited, or other BF neurons not directly inhibited, with the same neuronal order across heatmaps. Directly inhibited neurons were sorted by inhibition latency; the remaining BF bursting and other BF neurons were sorted by sound-response amplitude and baseline firing rate, respectively. Each row represents one neuron. **B**, Mean firing-rate changes during somatic inhibition on sound and catch trials. Gray points indicate mean firing-rate changes for individual neurons during the first 200 ms after light onset; colored symbols and error bars indicate mean ± s.e.m. across neurons within the three groups defined in **A**. **C,D**, Corresponding analyses during inhibition of VTA DA axon terminals in the BF. Conventions are as in **A,B**. Somatic and terminal inhibition significantly suppressed both cue-evoked and ongoing activity in directly inhibited BF neurons but did not significantly affect the other two groups.

**Figure S9.**
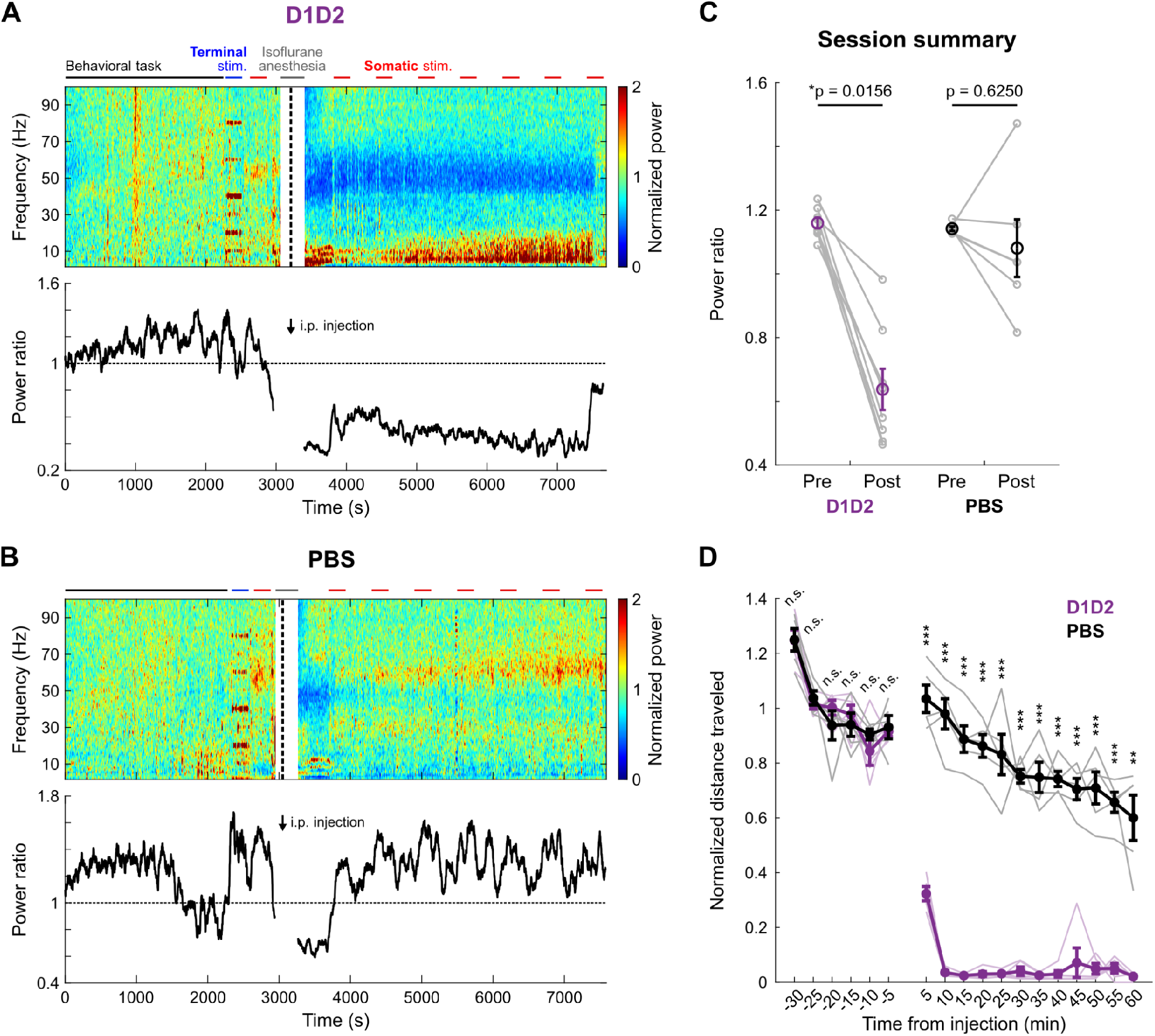
D1/D2 receptor blockade alters BF LFP power and locomotor activity. **A,B**, Example normalized BF LFP power spectrograms (top) and time-resolved high-to-low-frequency power ratios (bottom) before and after administration of D1/D2 receptor antagonists (**A**) or PBS (**B**). Arrows and vertical black lines indicate injection. Isoflurane and optogenetic stimulation epochs were excluded from analysis. **C**, Power ratios before and after injection in individual experiments. Gray lines indicate individual experiments; open black symbols show mean ± s.e.m. D1/D2 receptor blockade significantly reduced the power ratio, whereas PBS had no significant effect (two-sided Wilcoxon signed-rank tests with Bonferroni correction, *p < 0.05). **D,** Freely moving locomotor activity quantified as total distance traveled before and after injection. D1/D2 receptor blockade, but not PBS, significantly decreased voluntary movement (Two-way mixed-design repeated-measures ANOVA with Greenhouse–Geisser correction, with Bonferroni corrections for each comparison. **p < 0.01; ***p < 0.001).

**Table S1.** Summary of animals and sessions contributing to each experiment. Numbers, sex, and species of animals used in each experiment are listed alongside session counts. Sample sizes for spike and LFP analyses are reported separately.

| Experiment | Animal / Session | Figure |
| --- | --- | --- |
| Lick-schedule reward task | 7 male rats / 10 sessions | Fig. 1, 2, S2-S4 |
| Reward learning tasks | 3 male rats / 8 sessions | Fig. 3, S5 |
| DA somatic activation | 6 male, 1 female mice / 18 sessions | Fig. 4, S6 |
| DA terminal activation | 6 male, 1 female mice / 18 sessions | Fig. 4, S6 |
| DA somatic inhibition | 3 male, 2 female mice / 9 sessions | Fig. 5, S7 |
| DA terminal inhibition | 4 male, 1 female mice / 12 sessions | Fig. 5, S7 |
| DA somatic activation (D1/D2) | 3 male, 1 female mice / 10 sessions (spike)<br>2 male, 1 female mice / 8 sessions (LFP) | Fig. 6, S8 |
| DA somatic activation (PBS) | 3 male mice / 8 sessions (spike)<br>2 male, 1 female mice / 6 sessions (LFP) | Fig. 6, S8 |
| DA receptor antagonist testing (D1/D2) | 4 male, 1 female mice | Fig. S8 |
| DA receptor antagonist testing (PBS) | 5 male mice | Fig. S8 |
| ATLAS | 6 female mice | Fig. 6 |

## Acknowledgements

We thank T.-W. Chen for critical discussions of the manuscript; S.-T. Wu for technical support.

## Contributions

Y.-J.J. and S.-C.L. conceived and designed the study. Y.-J.J. conducted the main experiments and analyzed the data with S.-C.L. M.-C.C. and M.-E.K. performed the initial DA activation experiments. C.-Y.C. performed video-based tracking analysis. C.-Z.L. and A.S. developed the behavioral apparatus and instrumentation for head fixation and electrophysiological recordings. A.G. and F.G. performed the ATLAS tracing experiments. Y.-J.J. and S.-C.L. wrote the manuscript with input from all authors. S.-C.L. supervised the research.

## Corresponding author

Correspondence to Shih-Chieh Lin

